# Scoring of pathogenic non-coding variants in Mendelian diseases through supervised learning on ancient, recent and ongoing purifying selection signals in human

**DOI:** 10.1101/363903

**Authors:** Barthélémy Caron, Yufei Luo, Antonio Rausell

## Abstract

The study of rare Mendelian diseases through exome sequencing typically yields incomplete diagnostic rates, ~8-70% depending on the disease type. Whole genome sequencing of the unresolved cases allows addressing the hypothesis that causal variants could lay in non-coding regions with damaging regulatory consequences. The large amount of rare and singleton variants found in each individual genome requires computational filtering and scoring strategies to gain power in downstream statistical genetics tests. However, state-of-the-art methods estimating the functional relevance of non-coding genomic regions have been mostly characterized on sets of variants largely composed of trait-associated polymorphisms and associated to common diseases, yet with modest accuracy and strong positional biases. In this work we first curated a collection of n=737 high-confidence pathogenic non-coding single-nucleotide variants in proximal *cis*-regulatory genomic regions associated to monogenic Mendelian diseases. We then systematically evaluated the ability to predict causal variants of a comprehensive set of natural selection features extracted at three genomic levels: the affected position, the flanking region and the associated gene. In addition to inter-species conservation, a comprehensive set of recent and ongoing purifying selection signals in human was explored, allowing to capture potential constraints associated to recently acquired regulatory elements in the human lineage. A supervised learning approach using gradient tree boosting on such features reached a high predictive performance characterized by an area under the ROC curve = 0.84 and an area under the Precision-Recall curve = 0.47. The figures represent a relative improvement of >10% and >34% respectively upon the performance of current state-of-the-art methods for prioritizing non-coding variants. Performance was consistent under multiple configurations of the sets of variants used for learning and for independent testing. The supervised learning design allowed the assessment of newly seen non-coding variants overcoming gene and positional bias. The scores produced by the approach allow a more consistent weighting and aggregation of candidate pathogenic variants from diverse non-coding regions within and across genes in the context of statistical tests for rare variant association analysis.

## Introduction

To date, more than 4,000 Mendelian diseases have been clinically recognized ^1^, collectively affecting more than 25 million people in the US only ^2^. However, around 50% of all known Mendelian diseases still lack the identification of the causal gene or variant ^3^. Moreover, every year approximately 300 new Mendelian diseases are described, whereas the pace for discovery of the causal molecular mechanisms fluctuates at around 200 yearly ^3^. Despite the progress achieved through Whole Exome Sequencing (WES)-based studies, recent reviews show highly heterogeneous diagnostic rates across disease types ^4,5^, ranging from <15% (e.g. congenital diaphragmatic hernia or syndromic congenital heart disease) to >70% (e.g. ciliary dyskinesia). In those scenarios, a common working hypothesis is that non-coding variants could explain the etiology of many of the unresolved cases ^5^. Whole Genome Sequencing (WGS) permits to expand the survey of pathogenic variants to non-coding genomic regions in an unbiased way. Such possibility generates great expectations, as most trait/disease-associated Single Nucleotide Variants (SNVs) identified by Genome Wide Association Studies (GWAS) map to non-coding regions, suggesting a prominent role of regulatory elements in genetic diseases ^6,7^. Nevertheless, the large amount of rare and singleton variants in non-exonic positions shown by large-scale WGS projects in human ^8^, makes computational predictions a fundamental step to prioritize candidate variants for further clinical and experimental follow up.

A number of machine-learning methods have been developed in the last years to predict the regulatory consequences of non-coding SNVs ^9–15^. Two complementary perspectives have been exploited: First, from an evolutionary standpoint, genomic positions under non-neutral evolution are expected to have a functional role. Consequently, position-based purifying selection scores determined at different time-scales (i.e. from vertebrates, mammals and primates sequence alignments) have been successfully used by reference methods. Second, from a mechanistic view, phenotypic consequences of genetic variants are thought to result from their impact on non-coding functional elements, defined as those having reproducible biochemical features associated to regulatory roles, such as promoters, enhancers, silencers, repressors, etc. Thus, computational methods have exploited diverse sets of chromatin and epigenetic characteristics (e.g. histone marks, chromatin states, DNase I-hypersensitivity sites and transcription factor binding sites) obtained from heterogeneous sets of cell lines, primary cell types and tissues by Consortia such as ENCODE, FANTOM5, the Roadmap Epigenomics and BluePrint projects ^16–18^. While the ability of state-of-the-art methods to discriminate functionally relevant non-coding variants is well established, the value of such scores as a proxy of pathogenic potential in the context of Mendelian diseases is still unclear. This stems from the fact that functional scores of non-coding SNVs were mostly evaluated by their ability to identify trait-associated (e.g. quantitative trait loci, QTLs) and disease-associated loci from GWAS studies of common diseases. Yet, even in those contexts, predictive accuracy is modest and mainly driven by position-based interspecies conservation signals, with chromatin and epigenetic features providing only a marginal contribution ^10,13,15^. More recently, Smedley *et al.* developed the so-called Regulatory Mendelian Mutation Score (ReMM)^19^, which -to our knowledge-is the only method specifically developed to score pathogenic non-coding variants in the context of Mendelian disease studies. The approach trained a random forest classifier on a curated set of 406 SNVs (including long non-coding RNA SNVs). Twenty-six features were considered, including 8 interspecies conservation scores, 4 GC/CpG-based characteristics and 8 epigenetic features. Despite the simplicity of the model, ReMM scores proved valuable to prioritize Mendelian disease variants when integrated in a more comprehensive framework considering candidate regulatory regions and the phenotypic relevance of the associated genes ^19^.

In this work we hypothesized that the computational prediction of pathogenic non-coding variants in Mendelian diseases would benefit from a more comprehensive set of natural selection signals, notably regarding recent and ongoing selective constraints in human. In this regards, evolutionary and functional evidence support a rapid turnover of functional non-coding elements across species that would limit the capacity of interspecies conservation to pinpoint recently acquired regulatory sequences in the human lineage ^20,21^. Moreover, it has been suggested that lineage-specific and ongoing natural selection in human could help further understanding the partial overlap observed among the fraction of the genome inferred to be functional from evolutionary, biochemical and genetic evidences ^22,23^. The use of recent and ongoing purifying selection signals to prioritize pathogenic variants has been historically challenged by a number of confounding factors shaping patterns of human genomic variation. Thus, random genetic drift, population structure and demographic processes such as rapid expansions, migrations and population bottlenecks, have played a major role of governing changes in allele frequency within and between populations^23–25^. In addition, uneven recombination rates across the genome and heterogeneous neutral mutation rates^26^ associated to sequence context^27,28^ or to different types of non-coding elements^29^ further complicates the distinction of neutral versus non-neutral evolution.

Notwithstanding, the increasing sample size of current large-scale whole genome sequencing projects of the general population are providing a better resolution of recent and ongoing purifying selection signals in human^8,30–32^ that could improve their utility in scoring systems of pathogenic variants. In addition, machine learning methods have shown able to extract complex patterns associated to functional variants combining different types of selective constraints that would be missed by classical approaches^10,15^. Hence, a supervised learning approach could help better exploiting recent natural selection features in spite of confounding factors. To test both previous possibilities, in this study we first extracted a comprehensive set of recent and ongoing natural selection features determined from recent large-scale WGS projects in human together with interspecies conservation scores assessed on different evolutionary timescales. We then trained NCBoost, a classifier of non-coding SNVs based on gradient tree boosting, on a curated set of high-confidence pathogenic non-coding SNVs associated to monogenic Mendelian disease genes and on common non-coding SNVs without clinical assertions. The approach outperformed existing state-of-the-art methods under multiple training and testing scenarios, while overcoming gene and positional bias.

## Material and Methods

### High-confidence pathogenic variants

Three sets of high-confidence pathogenic variants in non-coding regions were obtained: 1) Regulatory disease-causing mutations (so-called “DM” set) from the Human Gene Mutation Database (HGMD, professional version, accessed on 2018/01/03, ^33^), manually annotated as involved in conferring the associated clinical phenotype; 2) pathogenic single-nucleotide variants (SNVs) from Clinvar ^34^ manually annotated as “pathogenic” with no conflicting assertions (GrCh37 release from 2017/12/31, downloaded from ftp://ftp.ncbi.nlm.nih.gov/pub/clinvar/); and 3) a manually curated set compiled from the medical literature of non-coding single-nucleotide variants associated with Mendelian disease and validated by experimentation or co-segregation studies, or for which other convincing evidence of pathogenicity was available (^19^.

### Variant mapping and annotation of non-coding SNVs

Only Single Nucleotide Variants (SNVs) where considered through the study. Variants were annotated using Annovar ^35^, downloaded on 2016-02-01; using the gene-based annotation option based on RefSeq for Humans (assembly version hg19); http://annovar.openbioinformatics.org/en/latest/user-guide/gene/) in order to obtain i) the gene region affected by intragenic variants, or ii) the nearest flanking gene in the case of intergenic variants. Exonic variants and variants within 10 base pairs (bp) of a splicing junction of protein-coding genes were removed (Annovar *splicing*_*threshold* =10). At this stage, variants from HGMD-DM, Clinvar and Smedley’2016 overlapping non-coding RNAs within an exon (n= 143, 2 and 68, respectively), intron (n= 24, 3, 13, respectively) or 10bp from a splicing junction (n= 1, 0, 0, respectively) were filtered out. In the case of SNV overlapping several types of regions associated to different genes or transcripts, the following three criteria were consecutively adopted: A) the default Annovar precedence rule for gene-based annotation was adopted, i.e.: exonic = splicing > ncRNA > UTR5 = UTR3 > intronic > upstream = downstream > intergenic. B) if after applying the previous precedence rule a SNV could still be associated to several neighbor/overlapping genes (e.g. in the intergenic region between two genes, or in the intronic region of two overlapping genes, etc), the SNV’s nearest protein coding gene was kept as a reference for the annotation of the variant. The SNV’s nearest gene was determined by the shortest distance to either the TSS or TSE. C) In case of SNVs with two or more genes with identical shortest distance to TSS/TSE, the SNV was tagged as ‘conflicted’ and filtered out from the analysis. After all previous filtering steps, a total of 18 disease-causing SNVs overlapping upstream (n=9), UTR’3 (n=7) and downstream regions (n=2) of non-coding RNAs were filtered-out. Thus, for the purpose of this study, the set of non-coding variants was constituted of SNVs associated to protein-coding genes and overlapping intronic, 5’ UTR or 3’ UTR, upstream and downstream regions -i.e. closer than 1kb from the Transcription Start Site (TSS) and the Transcription End Site (TSE) respectively- and intergenic regions.

### Curation of high-confidence pathogenic non-coding SNVs associated to monogenic Mendelian disease genes

For high-confidence pathogenic SNVs, we manually supervised a total of n=71 cases showing a disagreement between the gene associated to the variant in the original resource (i.e. HGMD-DM, Clinvar and Smedley’2016) and the gene associated by the previously described annotation procedure. The original gene assignment was kept for n=17 SNVs where conflict originated due to straightforward exceptions of the Annovar’s precedence rule or the assignment to the nearest upstream or downstream gene (Criteria A and B described in the previous section). The number of variants retained at this stage is represented in Figure 1A. Only high-confidence pathogenic non-coding variants associated to the same gene by both the original resource and the annotation process done in this work were retained for downstream analyses.

**Figure 1.**
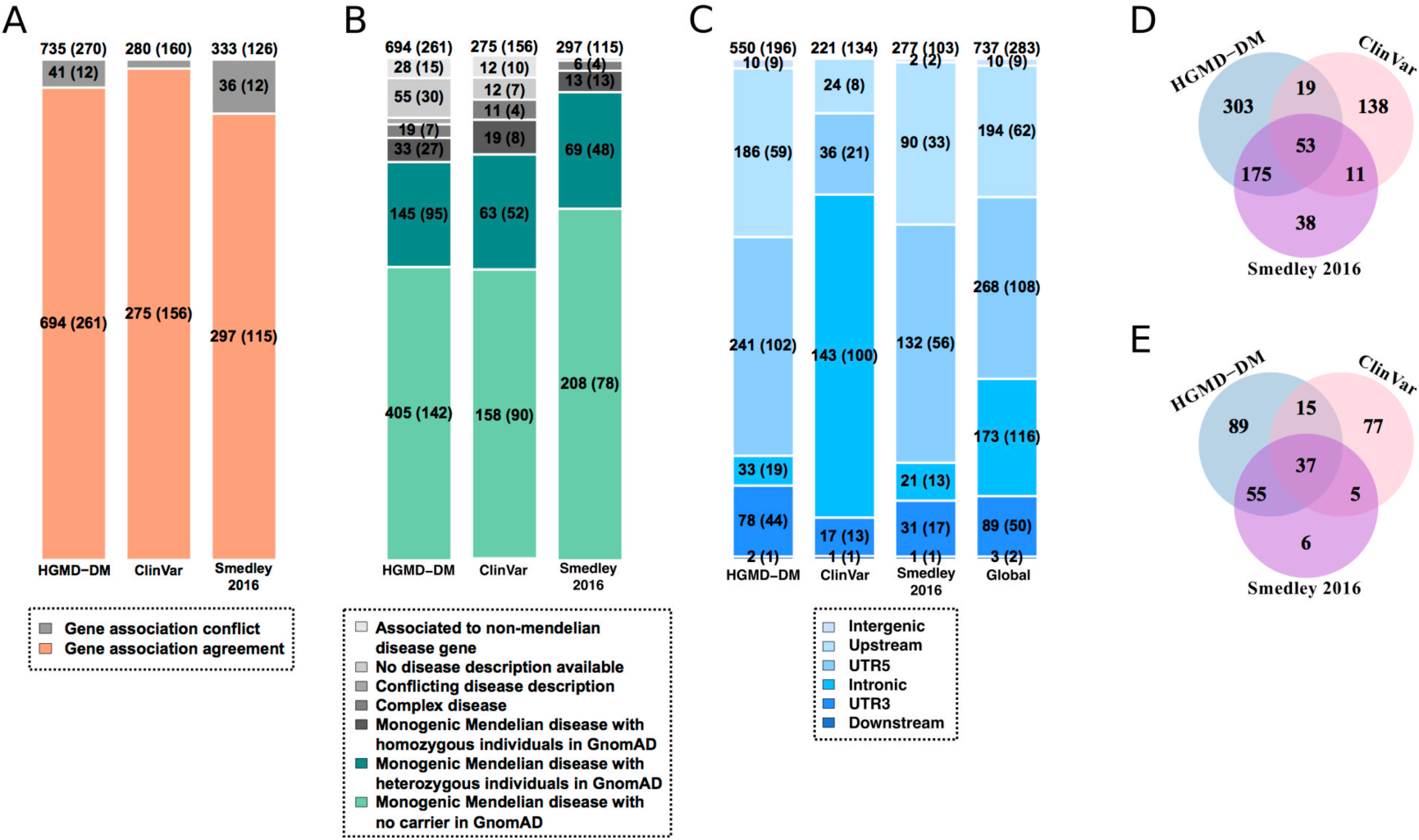
Curation of high-confidence pathogenic non-coding SNVs associated to monogenic Mendelian disease genes. **A.** Number of high-confidence pathogenic noncoding SNVs obtained from the Human Gene Mutation Database ^33^ (HGMD-DM), Clinvar ^34^ and Smedley’2016 ^19^ (**Methods**), after filtering out SNVs overlapping exonic and splice sites of protein-coding genes and exons and introns of non-coding RNAs. Only high-confidence pathogenic non-coding variants associated to the same protein-coding gene by both the original resource and the annotation process done in this work (depicted in orange), were retained for downstream analysis. **B.** Retained variants in (A) were further classified according the OMIM category of the associated gene, i.e.: non-Mendelian disease gene, Mendelian disease gene associated to a disease phenotype differing from the one reported in the original resource (i.e. presenting a conflicting disease description), complex Mendelian disease genes and monogenic Mendelian disease genes. Only high-confidence pathogenic non-coding SNVs associated to monogenic Mendelian diseases with no homozygous individuals in GnomAD database ^31^)depicted in green) were finally retained for downstream analyses. **C.** Distribution of the high-confidence pathogenic non-coding SNVs associated to monogenic Mendelian disease genes according to the type of gene region they overlap: intronic, 5’UTR, 3’UTR, upstream, downstream and intergenic regions. **D.** Distribution of the high-confidence pathogenic non-coding SNVs associated to monogenic Mendelian disease genes according to the original annotation source, i.e. HGMD-DM, Clinvar and Smedley’2016. **E**. Corresponding number of monogenic Mendelian disease genes collectively involved by SNVs in (D). The number of SNVs in each category is indicated inside the barplots and Venn diagrams together with the number of genes collectively involved (in parenthesis in A-C; totals are reported above each barplot).

We then evaluated whether the genes associated to high-confidence pathogenic non-coding SNVs were reported as Mendelian disease genes in ^1^. A list of n=3695 Mendelian disease genes was obtained following ^3^: OMIM raw data files mim2gene.txt, genemap2.txt and morbidmap.txt were downloaded from www.omim.org on 2017/10/13. MIM phenotype number and supporting evidence annotations where extracted from morbidmap.txt. Phenotype descriptions containing the word ‘somatic’ were flagged as ‘somatic’, those containing [‘carcinoma’, ‘cancer’, ‘tumor’, ‘leukemia’, ‘lymphoma’, ‘sarcoma’, ‘blastoma’, ‘adenoma’, ‘cytoma’, ‘myelodysplastic’, ‘Myelofibrosis’ or ‘oma’] were flagged as ‘cancer’, and those containing [‘risk’, ‘quantitative trait locus’, ‘QTL’, ‘{’, ‘[’ or ‘susceptibility to’] were flagged as ‘complex’. Phenotypes flagged as ‘somatic’ and ‘cancer’ were classified as ‘somatic cancer’. Mendelian genes were then defined as the genes having a supporting evidence level of 3 (i.e. the molecular basis of the disease is known) and not having a ‘somatic’ flag. Two main categories of Mendelian disease genes where defined: monogenic Mendelian disease genes (n=3354) and complex Mendelian disease genes (n=596), i.e. those presenting mutation risk factors, quantitative-trait loci (QTL) or contributing to susceptibility to multifactorial disorders or to susceptibility to infection ^3^. Of note, 255 genes were associated to both monogenic and complex Mendelian disease genes.

High-confidence pathogenic non-coding SNVs associated to monogenic Mendelian disease genes where manually further inspected to check consistency between the disease phenotype reported in the original source (i.e. HGMD-DM, Clinvar and Smedley’2016) and the ones described in OMIM database for the same gene. A total number of n=138 variants for which the agreement was unclear or a disagreement was observed were filtered out for downstream analyses. In the remaining set of high-confidence pathogenic non-coding SNVs associated to monogenic Mendelian disease genes, we then inspected whether variants were detected as heterozygous or homozygous among the individuals included in the GnomAD database ^31^; http://gnomad.broadinstitute.org/downloads, version r2.0.2, using both whole genome sequencing data and whole exome sequencing data. Variants present as homozygous in at least one carrier were filtered out for downstream analysis. Thus only high-confidence pathogenic non-coding SNVs associated to monogenic Mendelian diseases, with no homozygous individuals in GnomAD and overlapping intronic, 5’UTR, 3’UTR, upstream, downstream and intergenic regions were finally retained for downstream analysis (**Table S1**).

### Common and rare human variants without clinical assertions

Common and rare human variants without clinical assertions where obtained from dbSNP (downloaded on 2017/07/10 from ftp://ftp.ncbi.nih.gov/snp/organisms/human_9606/VCF/All_20170710.vcf.gz). For the purpose of this study, variants labeled as common (“COMMON=1”) and with Minor Allele Frequency (MAF)> 0.05 were considered as ‘common variants’, while those labeled as non-common (“COMMON=0”) and with MAF< 0.01 were annotated as ‘rare variants’. Variants with no MAF information (no “CAF” field reported in the “INFO” field of the variant) and multiallelic variants were filtered out. Common and rare human variants without clinical assertions were first annotated by Annovar and filtered as described above. For consistency in the comparison against the pathogenic set of SNVs, the set of common and rare human variants without clinical assertions was restricted to SNVs associated to protein-coding genes and overlapping intronic, 5’UTR, 3’UTR, upstream, downstream and intergenic regions. The list of protein-coding genes was extracted from Ensembl Biomart ^36^; human genome assembly version GrCh37.p13).

### Pathogenicity scores of non-coding SNVs

Pre-computed pathogenicity scores of non-coding SNVs were extracted from the following state-of-the-art methods: CADD non-coding score (version v1.3, ^9^; DeepSEA functional significance score (version v0.94,^13^; Eigen and Eigen-PC scores (version v1.1,^15^; FunSeq2 score (version v1.2, ^11^ and ReMM scores (version v0.3.1; ^19^).

### Feature extraction of non-coding SNVs

Features extracted are summarized in **Table 1**. They can be gathered in 5 main categories:

**Table 1.**
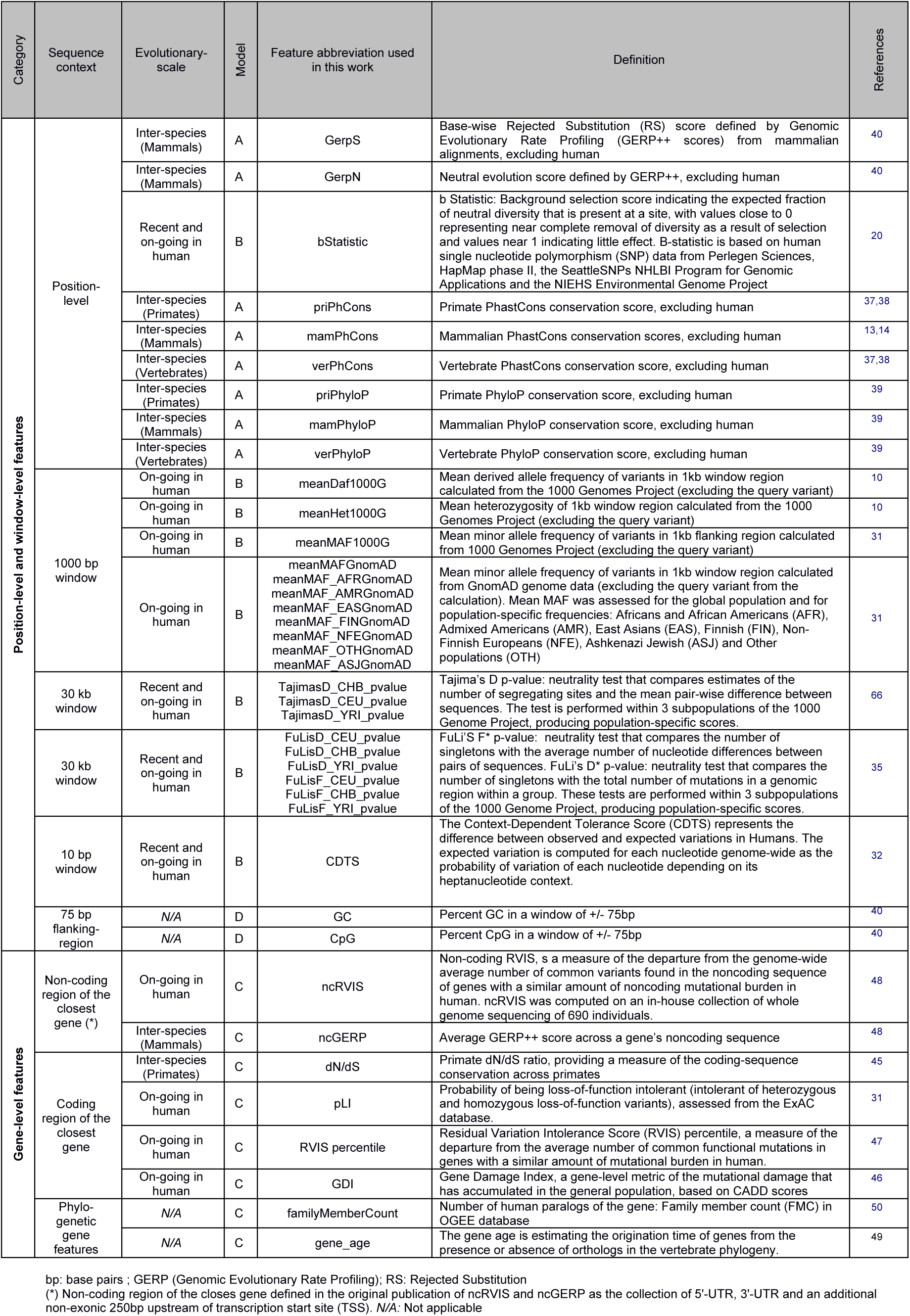
Natural selection features associated to non-coding Single-Nucleotide Variants mined in this work. Features are classified under different categories depending on the sequence context (i.e. position level, window level and gene level) and evolutionary scale: interspecies (vertebrates, mammals and primates, excluding human), or recent and ongoing natural selection in human. We note here that the query variant was excluded from the calculations involving mean allele frequencies and mean heterozygosity of a given region, and that the variant allele frequency itself was not used as a feature in any training or pathogenicity prediction throughout the study.

#### A. Inter-species sequence conservation features at position and window level

To evaluate evolutionary conservation at a given site, the following scores evaluating non-neutral rates of substitution from multiple species alignments (excluding human) were used: PhastCons ^37,38,39^ and PhyloP scores for three multi-species alignment (Vertebrates, Mammals and Primates, excluding human) and GerpN and GerpS single-nucleotide scores fom mammalian alignments ^40^, all of them obtained from CADD (version v1.3, ^*9*^, file name: whole_genome_SNVs_inclAnno.tsv.gz, downloaded from http://cadd.gs.washington.edu/download). PhyloP scores measure neutral evolution at individual sites. The score corresponds to the -log p-value of the null hypothesis of neutral evolution. Positive values (up to 3) represent purifying selection, while negative values (up to - 14) represent acceleration. PhastCons scores estimate the probability that the locus is contained in a conserved element. GerpN and GerpS single-nucleotide scores assess respectively the neutral substitution rate and the rejected substitution rate of the locus. A high GerpN value indicates high homology of the locus across species. Positive values of GerpS indicate a deficit in substitutions, while negative values convey a substitution surplus.

#### B. Recent and ongoing natural selection signals in Humans at position and window level

Three human population-specific natural selection scores based on the allele frequency spectrum on a 30 kb sequence window region centered around the SNV were obtained from *The 1000 Genomes Selection Browser 1.0* (http://hsb.upf.edu/,^41^): Tajima’s D ^42^, Fu & Li’s D^⋆^ and Fu & Li’s F* ^43^. Tajima’s D is a neutrality test comparing estimates of the number of segregating sites and the mean pair-wise difference between sequences. Fu & Li’s D* is a neutrality test comparing the number of singletons with the total number of nucleotide variants within a region. Fu & Li’s F is a neutrality test comparing the number of singletons with the average number of nucleotide differences between pairs of sequences. The three tests were performed within 3 populations of the 1000 Genome Project phase 1 data, producing population-specific scores: Yoruba in Ibadan, Nigeria (YRI), Han Chinese in Beijing, China (CHB) and Utah Residents with Northern and Western European Ancestry (CEU). Negative logarithmic percentiles associated to each of these score were used with values ranging from 0 (indicating positive selection) to 6 (indicating purifying selection). Here, we used the negative logarithm of the ranked percentile of each score over the whole genome (http://hsb.upf.edu/?page_id=594) associated to the raw scores, with values ranging from 0 (indicating positive selection) to 6 (indicating purifying selection).

The *background selection score* (B statistic, ^44^), indicating the expected fraction of neutral diversity that is present at a site, was obtained from CADD annotations (version v1.3). B statistic values close to 0 represent nearly complete removal of diversity as a result of selection and values near 1 indicate no conservation. B-statistic is based on human single nucleotide polymorphism (SNP) data from Perlegen Sciences, HapMap phase II, the SeattleSNPs NHLBI Program for Genomic Applications and the NIEHS Environmental Genome Project.

Context-dependent tolerance scores (CDTS) for 10bp bins of the human genome computed on 15496 unrelated individuals from the gnomAD consortium ^32^) were downloaded from http://www.hli-opendata.com/noncoding/ (file CDTS_diff_perc_coordsorted_gnomAD_N15496_hg19.bed.gz). The CDTS represents the difference between observed and expected variations in humans. The expected variation is computed genome-wide for each nucleotide as the probability of variation of each nucleotide depending on its heptanucleotide context. Low CDTS scores indicate loci intolerant to variation.

Mean heterozygosity and mean derived allele frequency of variants in a 1kb window region centered around the SNV and calculated from the 1000 Genomes Project (excluding the query variant) were obtained from ^10^ ftp://ftp.sanger.ac.uk/pub/resources/software/gwava/v1.0/source_data/1kg). Mean minor allele frequency (MAF) of variants in 1kb flregion were calculated from GnomAD genome data ^31^, excluding the query variant from the calculation. Mean MAF was assessed for the global population and for population-specific frequencies: Africans and African Americans (AFR), Admixed Americans (AMR), East Asians (EAS), Finnish (FIN), Non-Finnish Europeans (NFE), Ashkenazi Jewish (ASJ) and Other populations (OTH). Additionally, we extracted mean MAF of variants in a 1kb window calculated from the 1000 Genomes Project (excluding the query variant). MAFs both from GnomAD and the 1000 Genomes Project were extracted from GnomAD release r2.0.2, using the Genome VCF files (available at http://gnomad.broadinstitute.org/downloads).

#### C. Gene-based features

The following gene-level features associated to natural selection were obtained:

Primate dn/ds ratios (i.e. the ratio between the number of nonsynonymous substitutions and the number of synonymous substitutions) were taken from ^45^. Low dn/ds values reflect purifying selection, while high dn/ds values are indicative of positive selection.

The gene probability of loss-of-function intolerance (pLI), from ^31^, reflecting a depletion of rare and *de novo* protein-truncating variants as compared to the expectations drawn from a neutral model of de novo variation on ExAC exomes data. pLI values close to 1 reflect gene intolerant to heterozygous and homozygous loss-of-function mutations.

Gene Damage Index (GDI), a gene-level metric of the mutational damage that has accumulated in the general population, based on CADD scores ^46^. High GDI values reflect highly damaged genes.

The Residual Variation Intolerance Score (RVIS percentile; ^47^), which provides a gene measure of the departure from the average number of common functional mutations in genes with a similar amount of mutational burden in human. High RVIS percentiles reflect genes highly tolerant to variation.

The non-coding version of the RVIS score (ncRVIS, ^48^ measuring the departure from the genome-wide average of the number of common variants found in the noncoding sequence of genes with a similar amount of noncoding mutational burden in human. Negative values of ncRVIS indicate a conserved proximal non-coding region, while positive values indicate a higher burden of SNVs than expected under neutrality.

The average non-coding GERP (ncGERP) is the average GERP score ^40^ across a gene’s noncoding sequence (Petrovski *et al.*, 2015). Both in the case of ncRVIS and ncGERP, the non-coding sequence was defined in the original publication as the collection of 5’-UTR, 3’-UTR and an additional non-exonic 250 bp upstream of transcription start site (TSS).

Gene age estimating the gene time of origin based on the presence/absence of orthologs in the vertebrate phylogeny was taken from ^49^. It varies from 0 (oldest) to 12 (youngest, corresponding to human specific genes). The number of human paralogs for each gene was obtained from the OGEE database ^50^.

For all scores, gene names were mapped to approved gene symbols from HGNC. Missing values were imputed through the median value computed over all protein coding genes.

#### D. Sequence context

The percentage of GC and CpG in a window of 150bp around the variant of interest was taken from CADD v1.3 annotations. In addition, we one-hot encoded the non-coding genomic region overlapping the SNV annotated by AnnoVar, and used it as binary features: intronic, 5’UTR, 3’UTR, upstream, downstream and intergenic regions.

#### E. Epigenetic features

Epigenetic features such as histone modification marks, nucleosome position, open chromatin profiles and transcription factor binding sites (TFBS) profiles generated by the ENCODE project ^51^ were extracted from CADD v3.1 annotations. DNA accessibility was assessed using two set of features: 1) the open chromatin evidence coming from the open chromatin super track, containing peak signal and Phred-scaled p-values of evidence for five open chromatin assays: DNase-seq (EncOCDNaseSig and EncOCDNasePVal), FAIRE-seq (EncOCFaireSig and EncOCFairePVal) and ChIP-seq using CTCF (EncOCctcfSig and EncOCctcfPVal), PolII (EncOCpolIISig and EncOCpolIIPVal) and Myc (EncOCmycSig and EncOCmycPVal), the Phred-scaled combined p-value of both DNase-seq and FAIRE-seq assays (EncOCCombPVal) and the Open Chromatin Code (EncOCC), a metric integrating DNaseI, FAIRE and ChIP-seq peak evidence of open chromatin. Further details are provided herein: http://rohsdb.cmb.usc.edu/GBshape/cgi-bin/hgTrackUi?g=wgEncodeOpenChromSynth and http://rohsdb.cmb.usc.edu/GBshape/cgi-bin/hgTables?db=hg19&hgta_group=regulation&hgta_track=wgEncodeOpenChromSynth&hgt <u>a table=wgEncodeOpenChromSynthGm18507Pk&hgta doSchema=describe+table+schema</u>). And 2) the maximum nucleosome position score obtained through MNase-seq (EncNucleo), indicating packed chromatin states: http://genome.ucsc.edu/cgi-bin/hgTrackUi?db=hg19&g=wgEncodeSydhNsome. Potential transcription factor activity was assessed using i) the number of different overlapping TFBS (TFBS), ii) the number of overlapping TFBS peaks summed over cell types (TFBSPeaks) and iii) the highest value of overlapping TFBS peaks across cell types from ChIP-seq (TFBSPeaksMax), as well as using histone modification marks, such as the maximum methylation peak at H3K4 (EncH3K4Me1, enhancers-associated), maximum trimethylation peak at H3K4 (EncH3K4Me3, promoter-associated) and maximum acetylation peak at H3K27 (EncH3K27Ac, associated to active enhancers).

### NCBoost training strategy

NCBoost training was performed with XGBoost, a machine learning technique based on gradient tree boosting (aka. gradient boosted regression tree; ^52,53^). The R implementation from https://github.com/dmlc/xgboost (version 0.71.1) was used with parameters: eta=0.01, max_depth=25 and gamma=10, selected to avoid overfitting and after parameter optimization through a prior tenfold cross-validation step.

To train NCBoost, we first randomly split the complete list of protein-coding genes in 10 genome partitions of equal size, with the same distribution across all chromosomes and keeping in each of them the same proportion of monogenic Mendelian disease genes presenting high-confidence pathogenic non-coding variants (see above). Throughout the work, each disease-causing variant (aka. ‘positive’ variants) was associated with a unique set of 10 ‘negative’ variants, randomly sampled from the set of common human variants without clinical assertion described above and associated to genes within the same genome partition. Random sampling of common variants was matched to the positive set to keep the same fraction of variants per type of region: intronic, 5’UTR, 3’UTR, upstream, downstream and intergenic regions. A maximum of one positive and one negative variant associated to the same gene was allowed, although no minimum per gene was required. For the training step, a maximum of one disease-causing non-coding variant was randomly sampled per gene (**Table S2**). We then trained NCBoost as a bundle of 10 independently trained models, consecutively excluding in each of them 1 of the 10-genome partitions described above.

### Correlation between independently trained 10-NCBoost models

To assess the correlation among the scores led by the independently trained 10-NCBoost models, we created 11 genome partitions in order to create 11 independent sets of positive and negative variants, randomly sampled in an analogous way as described above. One partition was randomly selected and reserved for validation while the other 10 were used for training. 10-NCBoost models were then independently trained using the set of features A+B+C+D described above, by consecutively excluding in each of them 1 of the 10-genome partitions. Then, each model was used to score variants in the 11^th^ partition. Correlation among the scores of the 10 models was assessed through Spearman rank correlation.

### Random sampling of rare human variants without clinical assertion

Each disease-causing variant was associated with a unique set of 10 rare variants, randomly sampled from the set of rare human variants without clinical assertion described above. Random sampling of rare variants was matched to the positive set to keep the same fraction of variants per type of region: intronic, 5’UTR, 3’UTR, upstream, downstream and intergenic regions. A maximum of one positive and one rare variant associated to the same gene was allowed, although no minimum per gene was required.

### Region-based random sampling of common variants

To constitute a “region-context” matched set of positive and negative variants, each disease-causing variants was associated –when available-with one common variant, randomly sampled from the set of common human variants without clinical assertion associated to the same gene and mapping to the same region (intronic, 5’UTR, 3’UTR, upstream, downstream and intergenic regions). Disease-causing variants with no matching common variants in the same region of the same gene were excluded from the region-context matched set of positive and negative variants. Multiple positive-negative variant pairs per gene were allowed in this setting. In the end, 149 region-matched pairs of pathogenic and random common variants were sampled, associated to 54 unique genes.

### Annotation of dominant/recessive and haploinsufficient genes

A list of n=299 haploinsufficient genes was obtained from ^54^. Genes intolerant to heterozygous truncation (pLI>0.9; ^31^ were obtained from ExAC browser: file fordist_cleaned_exac_nonTCGA_z_pli_rec_null_data.txt downloaded from ftp://ftp.broadinstitute.org/pub/ExAC_release/release0.3.1/functional_gene_constraint/. Dominant and recessive disease gene predictions were obtained from DOMINO ^55^, file score_all_final_03.04.17.txt downloaded from https://wwwfbm.unil.ch/domino/download.html). Following Quinodoz *et al.*, 2017, DOMINO score, reflecting the predicted probability of a gene to harbor dominant changes, was used to establish five gene categories (i.e. recessive, likely recessive, rest, likely dominant, dominant), corresponding to the probability intervals <0.2, 0.2-0.4, 0.4-0.6, 0.6-0.8, >=0.8, respectively.

### Software availability

Scripts to annotate SNVs with all features used in this study, software to score pathogenicity with NCBoost (ABCD model) and genome-wide pre-computed scores will be available online at http:// [URL TBD] upon publication of the manuscript

## Results

### Curation of a high-confidence set of pathogenic non-coding variants associated to monogenic Mendelian disease genes

The number of high-confidence pathogenic non-coding variants obtained from HGMD-DM, Clinvar and Smedley’2016 is represented in **Figure 1A (Methods)**. The majority of causal variants were assigned to the closest protein-coding gene in the reference source (94%, 98% and 89%, respectively). Thus the available set is mostly constituted of proximal cis-regulatory variants (**Figure 1A**), with distal cis-regulatory and trans-acting variants scarcely represented. Our curation effort allowed further refining this set to retain the fraction of pathogenic variants confidently associated to monogenic Mendelian diseases genes (84%, 87% and 98%, respectively; **Figure 1B**). In addition, a small though non-negligible fraction of variants for which homozygous individuals were detected in recent large-scale whole exome and genome sequencing (GnomAD; ^17^) were excluded for downstream analysis (5%, 7% and 4% respectively; **Figure 1B**). After all filtering steps, a total of 737 pathogenic non-coding SNVs collectively associated to 283 genes were retained (**Figure 1C**). Variants distributed in intronic (23%), UTR’5 (36%) and UTR’3 (12%), and 1Kb-upstream TSS (26%), with a minority of variants in 1Kb-downtream TSE (<1%) and in intergenic regions (1%). The 3 resources mined in this work (HGMD-DM, Clinvar and Smedley’2016) showed varying degrees of overlap regarding causal SNVs (**Figure 1D**) and associated genes (**Figure 1E**). Notably, the set of 283 monogenic Mendelian disease genes collectively affected by pathogenic non-coding SNVs is enriched in haploinsufficient genes (Odds ratio OR = 2,59, one-sided Fisher test p-value= 1,279e-9), in genes intolerant to heterozygous truncation (OR = 1,29; p-value=1,279e-9) and in genes predicted to have a dominant inheritance mode (OR = 1,36; p-value=1,72e-3) as compared to a background set of 3354 monogenic Mendelian disease genes (**Figure S1**; **Methods**).**!**

### Distribution of state-of-the-art pathogenicity scores across pathogenic and non-pathogenic SNVs

We then checked the distribution of six state-of-the-art pathogenicity scores (CADD, DeepSea, Eigen, Eigen-PC, FunSeq2 and ReMM; **Methods**) across the 737 high-confidence non-coding pathogenic variants and 4’960’178 common SNVs without clinical assertions. All evaluated scores showed marked differences depending on the type of gene region involved (i.e. intergenic, intronic, 3’UTR, 5’UTR, upstream and downstream regions of associated genes; **Figure S2**). Thus, the distributions of median scores per gene for pathogenic SNVs in 5’UTR and for SNVs within 1Kb-upstream TSS were shifted towards more severe values than those of pathogenic SNVs in 3’UTR, intronic and intergenic regions. Bias per gene region was also observed across common SNVs without clinical assertions, suggesting that the regulatory region where a variant maps systematically biases the scores (**Figure S2**). Surprisingly, the distributions of median scores per gene for common SNVs in 5’UTR was not significantly lower (i.e. less severe) than that of pathogenic SNVs in 3’UTR, intronic and intergenic regions for none of the 5 scores evaluated (two-sided Wilcoxon test p-values for all pair-wise comparisons evaluated are reported in **Table S3**). As a corollary, the previous observations warn about the necessity of matching the relative composition of pathogenic and non-pathogenic SNVs across different gene regions in predictive benchmarks, as well as for relative differences in region distribution across datasets (**Figure 1C**).

### Ability of natural selection signals to predict pathogenic non-coding SNVs when considered independently

**Table 1** summarizes the set of natural selection features extracted for both pathogenic non-coding SNVs and common SNVs without clinical assertions. Features gathered covered different evolutionary scales and can be classified as interspecies natural selection (considering vertebrates, mammals and primates, excluding human) or recent and ongoing natural selection in human. Second, features were categorized either as “position-based”, when they refer to the specific genomic position where the variant occurred, “window-level” when they refer to a given sequence interval centered in the SNVs, or “gene-level”, when they refer to the global characteristics of the closest protein-coding gene.

For the purpose of this work, features were group under three main sets (**Table 1**): A) interspecies sequence conservation features at position and window level; B) recent and ongoing natural selection signals in human at position and window level; and C) gene-based features. Furthermore, we included 2 additional sets of features: D) the sequence context, i.e. GC and CpG content as well as information of the type of gene region (intronic, 5’UTR, 3’UTR, upstream, downstream and intergenic region); and E) epigenetic features such as histone modification marks, nucleosome position, open chromatin profiles and transcription factor binding sites (TFBS) profiles generated by the ENCODE project ^51^.

We first checked the predictive ability of each individual feature to classify the n=737 high-confident set of pathogenic non-coding SNVs associated to monogenic Mendelian disease genes from a ‘negative set’ of n=7370 randomly sampled common SNVs without clinical assertions and matched by region (**Methods**). **Figure 2** shows the area under the receiver operating characteristic (AUROC) curve and the area under the Precision-Recall curve (AUPRC) obtained for each feature. The ranking of features according to both AUROC and AUPRC showed that predictive ability was dominated by interspecies sequence conservation features at position and window level (Category A), while only poor performances were observed for the rest of features when considered independently.

**Figure 2.**
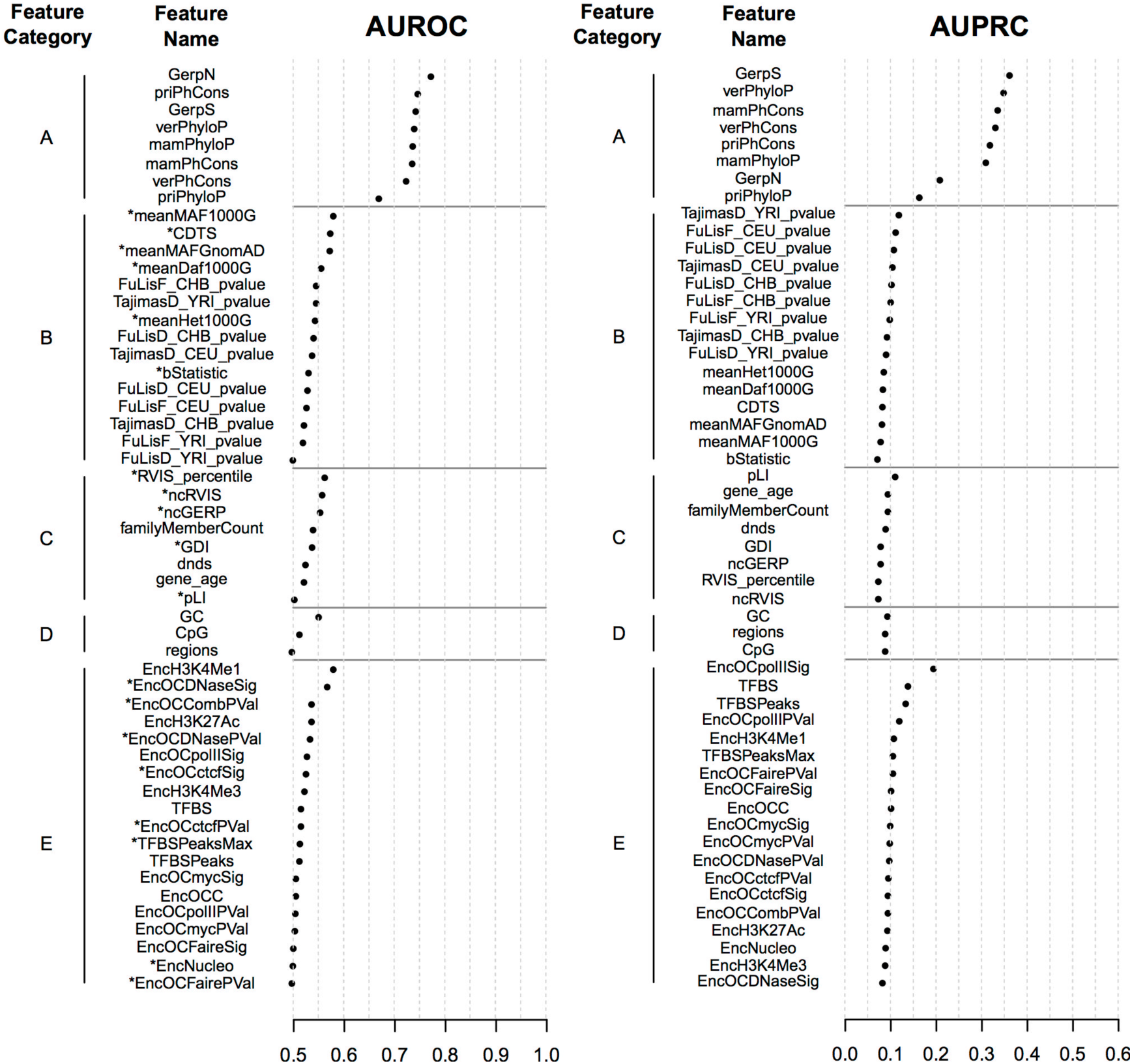
Performance of individual features mined in this work to classify high-confident set of pathogenic non-coding SNVs associated to monogenic Mendelian disease genes (n=737) from a ‘negative set’ of randomly sampled common SNVs without clinical assertions (n=7370) matched by region. The area under the receiver operating characteristic curve (AUROC; left panel) and the area under the Precision-Recall curve (AUPRC; right panel) obtained for each feature is represented. Features are gathered according to 5 categories (A-E; **Methods**), and ranked within category by decreasing AUROC and AUPRC. AUROC values <0,5 (anti-classifiers) were transformed in 1-AUROC values for the purpose of this figure and are indicated with an asterisk (^⋆^). Of note, population-specific GnomAD MAFs (**Methods**) are not shown for simplicity. 1 hot-encoded SNV region features (i.e. ‘intronic’, ‘UTR5’, ‘UTR3’, ‘upstream’, ‘downstream’ and ‘intergenic’) are gathered as a single feature labeled as ‘region’.

### Supervised learning of NCBoost based on a comprehensive set of ancient, recent and ongoing purifying selection signals in human

NCBoost, a machine learning approach based on gradient tree boosting (**Methods**) was trained on a ‘positive set’ of n=283 high-confident set of pathogenic non-coding SNVs associated to monogenic Mendelian disease genes (randomly selecting one variant per gene out of the total n=737 initially obtained to avoid gene-level contamination of the training/testing sets; **Figure 1C**) and a ‘negative set’ of n=2830 randomly sampled common SNVs without clinical assertions, matched by region and allowing a maximum of one negative variant per gene (**Methods**). NCBoost is a bundle of 10 independently trained models, consecutively excluding in each of them 1 out of 10 genome partitions were ‘positive’ and ‘negative’ variants are evenly distributed. In such a way, each non-coding variant in a putative cis-regulatory region of a protein-coding gene may be scored in NCBoost by the model that excluded from its training all variants -either pathogenic or non-pathogenic-associated to the same gene. This strategy permits to reduce overfitting as well as to avoid biasing the score of newly seen variants by the fact that they mapped in the vicinity of variants and genes initially presented to the classifier. Therefore, NCBoost may be applied to score any set of non-coding variants in cis-regulatory regions with no contamination with the training set. Of note, the 10 models proved to be largely equivalent among them, as shown by the high correlation of their scores when applied to an independent set of variants excluded from their training (average Spearman correlation 0.96 ± 0.0111 of all pairwise comparisons among the 10 models; **Methods**).

Six feature configurations were evaluated, including the following combinations of feature categories: A, B, A+B, A+B+C, A+B+C+D and A+B+C+D+E. The different NCBoost configurations were first tested mimicking a ten-fold cross-validation on the same n=283 high-confidence pathogenic non-coding SNVs and n=2830 common variants. **Figure 3** shows the area under the receiver operating characteristic (AUROC) curve and the area under the Precision-Recall curve (AUPRC) obtained for each of the six feature configurations. Best performance was reached by the model including ABCD features: AUROC_ABCD_ = 0,84 and AUPRC_ABCD_=0,47. The figures represent a relative improvement of 9% (AUROC) and 42% (AUPRC) over a model based purely in interspecies sequence conservation features at position and window level. Results were consistent when NCBoost was trained and tested on positive variants from each of the 3 resources taken independently, i.e. HGMD-DM, Clinvar and Smedley’2016 (**Figure S3A**, **S3B** and **S3C**, respectively).

**Figure 3.**
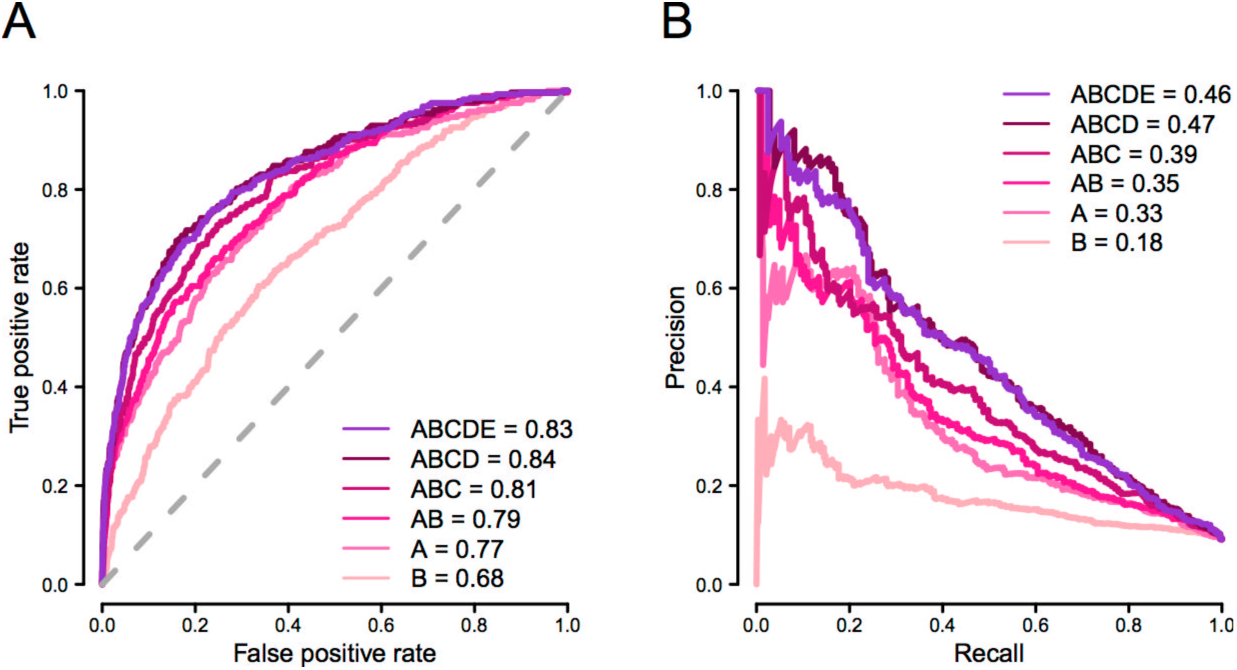
Comparative performance of NCboost models trained upon different sets of features. The figure represents the area under the receiver operating characteristic curve (AUROC; **Panel A**) and the area under the Precision-Recall curve (AUPRC; **Panel B**) obtained for each of the six feature configurations evaluated (feature categories A, B, A+B, A+B+C, A+B+C+D and A+B+C+D+E) when tested mimicking a ten-fold cross-validation on n=283 high-confidence pathogenic non-coding SNVs and n=2830 common variants without clinical assertions.

The features importance analysis of NCBoost upon the explored feature configurations, revealed a balanced contribution of inter-species sequence conservation features at position and window level (Category A, cumulative importance in the ABCD configuration CI_ABCD_= 42%) and recent and ongoing natural selection signals in human at position and window level (Category B, CI_ABCD_= 33%) collectively considered (**Figure S4A** and **S4B**). Such balance is observed in spite of the sharp differences in predictive ability observed across features when considered independently (**Figure 2**). The collective feature importance of recent and ongoing natural selection signals in human is in turn much higher than what could be expected from the observed incremental performance obtained in the join model AB (interspecies and intraspecies selection) as compared to B (interspecies selection; **Figure 3**). Both previous observations are not merely the straightforward consequence of the correlation structure across features (**Figure S5**). The previous results show the capacity of a supervised learning approach using a regression tree to extract complex patterns of natural selection signals distinguishing pathogenic versus non-pathogenic non-coding variants.

Notably, in contrast with the state-of-the-art methods evaluated (**Figure S2**), the per-region distribution of NCBoost scores across the 737 high-confidence non-coding pathogenic variants and 4’960’178 common SNVs, showed a clearer separation of between pathogenic and common variants for all types of regions evaluated (**Figure S6**). Thus, the distributions of median scores per gene for common SNVs in 5’UTR was significantly lower (i.e. less severe) than that of pathogenic SNVs in all evaluated regions (i.e. intronic, 3’UTR, 5’UTR and upstream regions; two-sided Wilcoxon test p-values <1e-10; **Table S3**), with the exception of intergenic region due to the low sample size.

### Comparative benchmark against state-of-the-art methods

NCBoost performance observed in **Figure 3** (Configuration ABCD) was compared against the results of the 6 state-of-the-art methods considered in this work (i.e. CADD, DeepSEA, Eigen, Eigen-PC, FunSeq2 and ReMM) when applied on the same ‘positive’ and ‘negative set’ of SNVs (**Figure 4**). NCBoost outperformed all evaluated methods both regarding AUROC and AUPRC, with a relative improvement of 10% and 34% respectively over the 2^nd^ ranked method (REMM), and of 13% and 104% over the 3^rd^ ranked method (Eigen). We note here that REMM is a supervised learning method whose training set partially overlapped with the ‘positive set’ of pathogenic non-coding SNVs variants used for testing here. Figures were consistent when the benchmark was performed on positive variants from each of the 3 resources taken independently, i.e. HGMD-DM, Clinvar and Smedley’2016 (**Figure S7A**, **S7B** and **S7C**, respectively).

**Figure 4:**
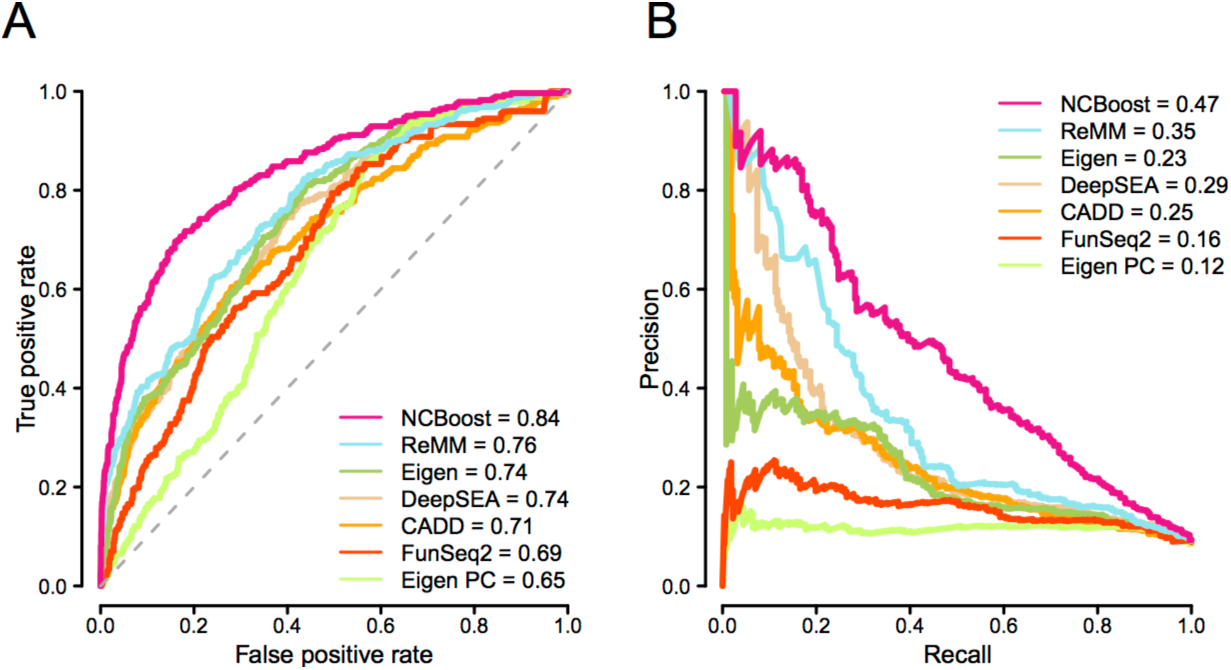
Comparative performance of NCBoost against state-of-the-art methods. Figure shows the represents the area under AUROC (**Panel A**) and the AUPRC (**Panel B**) obtained for NCBoost (configuration of features ABCD) together with 6 state-of-the-art methods (CADD, DeepSEA, Eigen, Eigen-PC, FunSeq2 and ReMM; **Methods**) when tested on the same set of ‘positive’ and ‘negative’ variants described for **Figure 3**.

The outperformance of NCBoost over reference methods was also observed when testing on the same ‘positive set’ of n=283 high-confident set of pathogenic non-coding SNVs as in **Figure 4** and on a negative set that -rather than of common variants-is composed of 2830 randomly selected rare variants (allele frequency < 1%) matched by region (**Figure S8**; **Methods**). This test allows ruling out the possibility that figures obtained in **Figure 4** are merely explained by the capacity to discriminate rare from common variants, rather than pathogenic from non-pathogenic variants.

In a more stringent set-up, we further explored the capacity of the different methods to discriminate pathogenic and non-pathogenic variants within the same non-coding region of a given gene. For this purpose, we restricted the previous testing to a set of 149 region-matched pairs of pathogenic and random common variants associated to 54 unique genes (**Figure S9**). Figures obtained were consistent with those previously observed in **Figure 4** and **Figure S8**, further supporting the superior capacity of NCBoost to discriminate pathogenic variants as compared to reference methods. We note that both in **Figure S8** and **Figure S9**, no re-training of NCBoost was done, but used the same NCBoost ABCD model trained as described in the previous section.

### Fully independent training and testing across all possible configurations of the three sources of high-confident non-coding pathogenic SNVs

To further characterize the performance of the NCBoost approach upon different training and testing scenarios, we evaluated all possible configurations of the training and testing set upon the three sources of high-confident non-coding pathogenic SNVs, i.e: HGMD-DM, Clinvar and Smedley’2016. Thus, the ‘positive set’ of n=283 high-confident set of pathogenic non-coding SNVs and the associated ‘negative set’ of n=2830 common SNVs matched by region (**Methods**) were each split in two non-overlapping sets in three different ways according to the source of annotation (**Table 2**), that is: training on the pathogenic variants reported in one source and testing on those in the other two sources not overlapping with the first one. In addition, we explored two additional configurations: training on variants reported in at least two sources and testing on those reported only in one single source, and the other way around. For each different training set, we retrained NCBoost as a bundle of 10 independently trained models, consecutively excluding in each of them 1 out of 10 genome partitions as previously done for the entire sets. We note again that a maximum of one positive and one negative variant per gene was allowed within the positive and negative sets, so that gene-level contamination across the training and test sets is avoided. **Table 2** shows the AUROC and AUPRC values obtained on each of the independent test sets when applying the corresponding NCBoost model. Consistently with previous sections, NCBoost outperformed the reference state-of-the-art methods under all training and testing scenarios evaluated.

**Table 2.** AUROC and AUPRC values obtained by NCBoost upon different configurations of the training and independent test sets. The figures obtained by the 5 state-of-the-art methods evaluated on the same test sets is shown together with NCboost.

| Positive set used for NCBoost training |  | Positive set used for testing |  | AUROC |  |  |  |  |  | AUPRC |  |  |  |  |  |
| --- | --- | --- | --- | --- | --- | --- | --- | --- | --- | --- | --- | --- | --- | --- | --- |
| Source | # SNVs | Source | # SNVs | DeepSEA | CADD | Eigen | FunSeq2 | ReMM | NCBoost | DeepSEA | CADD | Eigen | FunSeq2 | ReMM | NCBoost |
| HGMD-DM | 186 | ! HGMD-DM | 97 | 0,76 | 0,67 | 0,78 | 0,67 | 0,74 | **0,82** | 0,24 | 0,16 | 0,23 | 0,15 | 0,26 | **0,36** |
| CV | 107 | !CV | 176 | 0,74 | 0,72 | 0,73 | 0,68 | 0,77 | **0,78** | 0,31 | 0,27 | 0,24 | 0,15 | 0,37 | **0,38** |
| Smedley | 78 | !Smedley | 205 | 0,72 | 0,69 | 0,73 | 0,66 | 0,75 | **0,78** | 0,25 | 0,24 | 0,21 | 0,14 | 0,31 | **0,36** |
| >= 2 sources | 73 | 1 source | 210 | 0,72 | 0,68 | 0,73 | 0,66 | 0,75 | **0,78** | 0,24 | 0,23 | 0,21 | 0,14 | 0,31 | **0,31** |
| 1 source | 210 | >= 2 sources | 73 | 0,81 | 0,8 | 0,82 | 0,77 | 0,82 | **0,85** | 0,43 | 0,31 | 0,29 | 0,24 | 0,48 | **0,52** |

## Discussion

In this work we implemented a supervised learning approach, so called NCBoost, to classify pathogenic SNVs based on a comprehensive set of features at the position, flanking region and gene level, associated to interspecies, recent and ongoing selection in human. When trained and tested on multiple configurations of high confidence sets of pathogenic non-coding SNVs associated to monogenic Mendelian disease genes, the approach showed superior performance than reference methods. Notable improvements were observed on precision-recall rates. The context-specific assessment of natural selection signals permitted to overcome the pervasive regional bias observed in all evaluated reference methods, which e.g. tend to provide scores to non-pathogenic SNVs in 5’UTR not significantly different from the scores assigned to high-confident pathogenic SNVs in 3’UTR and intronic regions.

The curation process showed that current sets of high confidence large-effect pathogenic non-coding SNVs associated to monogenic Mendelian diseases are mostly constituted of proximal cis-regulatory variants associated to the closest protein-coding gene, in line with previous reports^19^. Such distribution most probably reflects a historical ascertainment bias towards such regions in previously described_ Mendelian genes, which is expected to be steadily overcome by unbiased WGS approaches ^5^. However, in the time being, the current status posses limits to the supervised learning and benchmark on distal *cis*- and *trans*-acting pathogenic regulatory variants with clinical implications in Mendelian diseases, and warns about the applicability of our approach and reference methods in such scope.

The approach implemented allowed us to evaluate the ability to prioritize pathogenic non-coding SNVs of recent and ongoing natural selection features in human when considered independently, collectively and in combination with interspecies conservation. While none of the features evaluated showed individual predictive strength (**Figure 2**), supervised learning performed through gradient tree boosting found complex patterns associated to pathogenic SNVs, reaching a significant performance combining multiple features (**Figure 3**). Detailed feature importance analysis showed a prominent contribution of recent and ongoing natural selection signals under all feature configurations evaluated. However, their final impact in the global performance of the classifier, while significant, is attenuated by the fact that some signals may be redundant with selective constrains already accounted for by interspecies conservation. Best figures were, nevertheless, obtained when the collective assessment of interspecies and intraspecies natural selection features was performed taking into consideration the sequence context where SNVs occurred, as informed by the selective signals accumulated by the associated gene and by the type of non-coding element involved.

This work represents a proof-of-concept of the added value of incorporating a large and heterogeneous set of recent and ongoing natural selection features under a supervised machine learning approach for the detection of pathogenic non-coding SNVs associated to Mendelian diseases. The rapidly increasing sample size of current large-scale WGS projects in the general population is expected to have a major impact in the capacity to detect additional and more accurate recent and ongoing natural selection signals in human, with a consequent repercussion in their use to identify pathogenic non-coding variants, as recently illustrated ^32,56,57^

In the last years, different large-scale projects have identified an important fraction of regulatory elements of the human genome, and the epigenetic insights are proving valuable to understand the functional consequences of disease-associated variants in those regions ^16–18^. However, in the setting of this work, the small set of epigenetic features evaluated had only a minor contribution to the classification of pathogenic SNVs associated to Mendelian diseases, in line with the results of previous analysis ^10,13,15^. On the one hand, this may suggest that the epigenetic signals evaluated here are partially redundant with natural selection features; a more exhaustive extraction of epigenetic features is however beyond the scope of this work. On the other hand, it may reflect a lack of specificity in regards of the cell types and tissues relevant for the heterogeneous set of Mendelian diseases considered here. In this line, recent studies are consolidating a view of regulatory mechanisms that is highly cell type-specific, where gene expression, DNA methylation, histone modifications, promoter interaction networks and transcription factor binding sites may substantially vary across tissues and developmental stages ^58–61^. Thus, the assessment of non-coding variants in the context of Mendelian diseases may largely benefit from the integration of purifying selection signals with the epigenetic information derived on the particular cell types, tissues and/or developmental time relevant for the onset and progression of a disease, as illustrated by recent successful examples ^57,62^. Notwithstanding, the identification of the specific cell type and tissue to be considered may be a challenging task, especially in the case of largely uncharacterized rare Mendelian diseases and syndromes.

Recently, it was shown that the number of singleton variants found on each newly sequenced genome stabilizes on average at ~8’500, with regulatory elements highly enriched in the relative amount of SNVs found per kb of sequence ^8^. The large amount of rare variants in each individual genome, together with the typically low number of participants in the study of specific rare diseases, challenges the statistical power of downstream statistical association and/or linkage studies to associate a genotype with a phenotype. The scoring approaches evaluated in this work may help filtering variants to increase power, although they often need to be integrated within more comprehensive frameworks in order to reach the necessary sensitivity and specificity to identify causal variants in disease cohorts^19^. In addition to the use of epigenetic information previously discussed, variant filtering strategies include focusing on SNVs associated to genes of phenotypic relevance for the disease under consideration^19,63^. From a complementary perspective, gene-based or region-based aggregation tests of multiple variants (a class of rare variant association tests) have been developed to evaluate cumulative effects of multiple genetic variants in a gene or region, with the aim of increasing power when multiple variants are associated with a disease ^64^, e.g. burden tests and variance component tests implemented in popular software such as PLINK/SEQ and SKAT. In these approaches, a continuous weight function can be used in the aggregation of rare variants in order to up-weight those predicted to have more damaging consequences. A similar weighting strategy can be proposed for rare-variant extensions of the Transmission Disequilibrium Test in the analysis of parent-child trio data ^65^. In both previous families of statistical tests for rare variant analysis of WGS from Mendelian diseases studies, the pathogenic scores led by the supervised learning approach implemented in this work, NCBoost, may be used to weight the aggregation of candidate pathogenic SNVs across heterogeneous cis-regulatory elements in a consistent way.

## Supplemental Data

Supplemental Data include nine figures and three tables and can be found with this article online.

## Acknowledgements

This work was supported by the French National Research Agency (ANR) grant ANR-17-RHUS-0002 - C’IL-LICO project of the second “Investissements ďAvenir” program and by the MSDAvenir fund, Devo-Decode project.

## Supplemental Table and Figure Legends

**Table S1. High-confidence pathogenic non-coding variants associated to monogenic Mendelian disease genes.**

**Table S2. Variants randomly sampled from the set of common human SNVs without clinical assertion associated to protein-coding genes used as the non-pathogenic set for training and testing in this work.** Sampling of variants was done to match the relative distribution across gene regions of the high-confidence pathogenic non-coding variants reported in **Table S1**.

**Table S3.** Regional bias of pathogenic score distributions. Two-sided Wilcoxon test p-values evaluating the null hypothesis that the per-gene median pathogenicity score distribution in 5’UTR for common non-coding SNVs is not different than the corresponding distribution for pathogenic variants in the 6 types of genomic regions evaluated, i.e: intergenic, intronic, 3’UTR, 5’UTR, upstream and downstream. The reported p-values are associated to the distributions represented in **Figure S2**.

|  | Common SNVs |  |  |  |  | Pathogenic SNVs |  |  |  |  |
| --- | --- | --- | --- | --- | --- | --- | --- | --- | --- | --- |
|  | Intergenic | Intronic | Downstream | UTR3 | upstream | Intergenic | Intronic | Downstream | UTR3 | Upstream |
| CADD | <1E-16 | <1E-16 | <1E-16 | <1E-16 | <1E-16 | 0,125 | 3,59E-03 | 0,814 | 1,46E-10 | 6,21E-06 |
| FunSeq2 | <1E-16 | <1E-16 | <1E-16 | <1E-16 | <1E-16 | 0,0112 | 0,014 | 0,436 | 1,96E-07 | 4,98E-03 |
| DeepSEA | <1E-16 | <1E-16 | <1E-16 | <1E-16 | <1E-16 | 0,301 | 3,16E-05 | 0,427 | 9,78E-12 | 1,60E-07 |
| Eigen | <1E-16 | <1E-16 | <1E-16 | <1E-16 | <1E-16 | 0,175 | 4,09E-05 | 0,685 | 7,94E-14 | 1,02E-04 |
| ReMM | <1E-16 | <1E-16 | <1E-16 | <1E-16 | <1E-16 | 0,919 | 3,49E-02 | 0,391 | <1E-16 | 1,73E-09 |
| NCBoost | <1E-16 | <1E-16 | <1E-16 | 4,98E-10 | <1E-16 | 0,767 | <1E-16 | 9,39E-11 | <1E-16 | <1E-16 |

**Figure S1.**
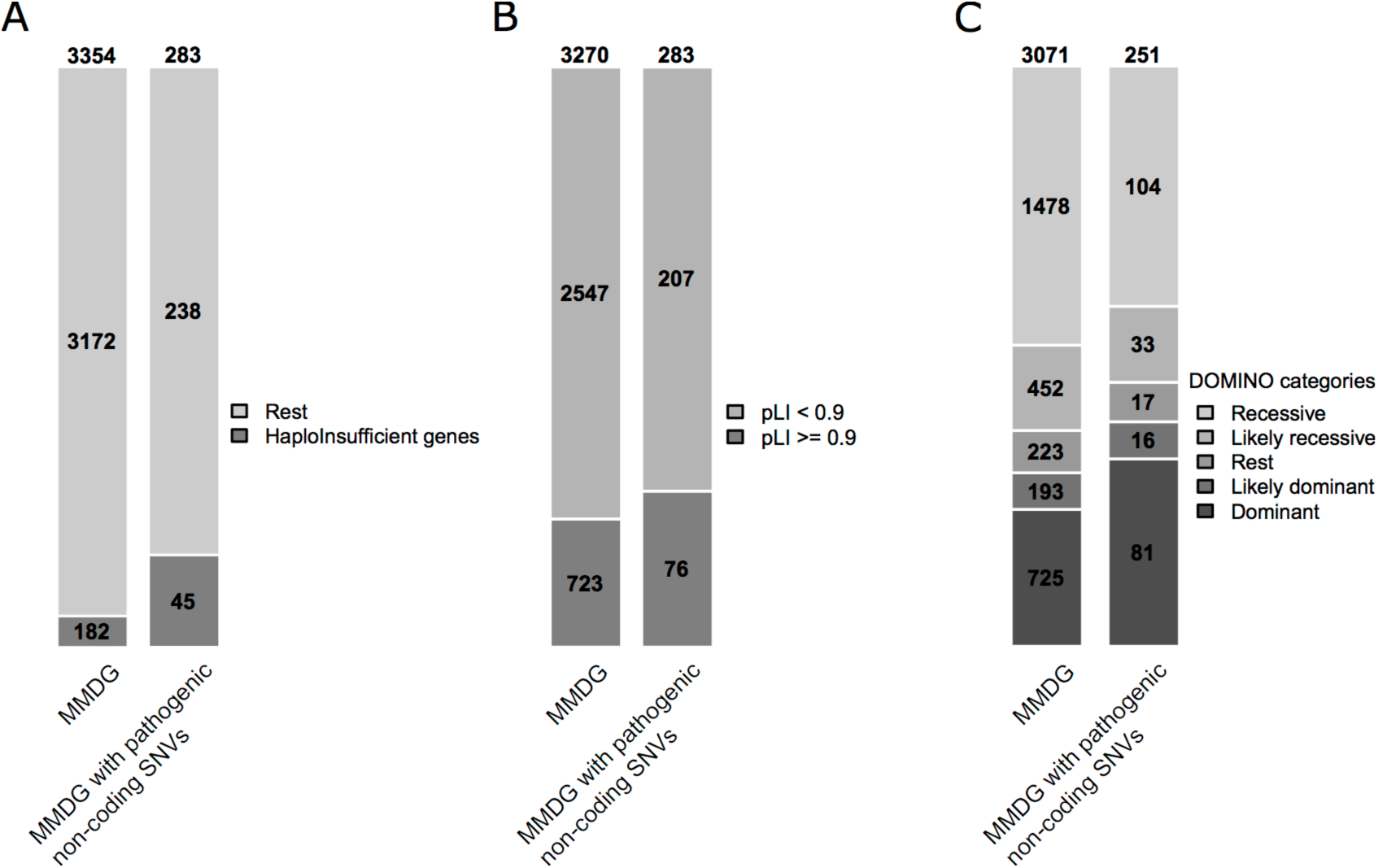
Relative enrichment of monogenic Mendelian disease genes associated to high-confidence pathogenic non-coding SNVs in haploinsufficient and dominant genes. Barplots showing the distribution of all monogenic Mendelian disease genes (n=3354; abbreviated as MMDG for the purpose of this figure) and those MMDG associated to high-confidence pathogenic non-coding SNVs (n=283) among the following categories: **A**. Haploinsufficient genes from ^54^. **B.** Genes intolerant to heterozygous truncation (pLI>0.9 ^31^). **C.** DOMINO gene predictions: recessive, likely recessive, rest, likely dominant, dominant (**Methods**). One-sided Fisher test p-values assessing the enrichment of monogenic Mendelian disease genes associated to high-confidence pathogenic non-coding SNVs in haploinsufficient and dominant genes were: 1.279e-09 (Panel A); 0.04107 (Panel B) and 0.00172 (Panel C).

**Figure S2.**
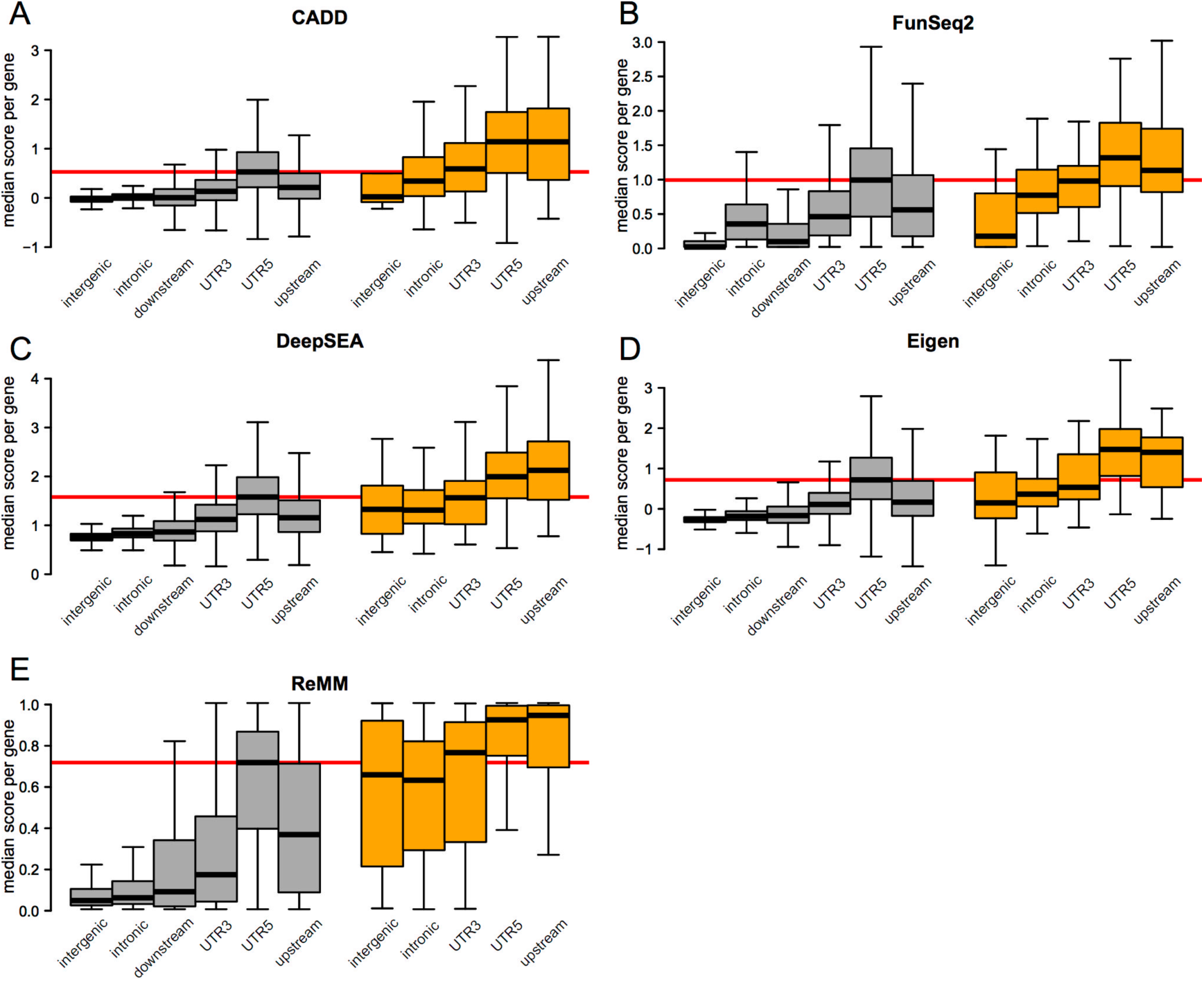
Distribution of pathogenic scores of non-coding SNVs according to the affected type of genomic region. Boxplots in the panels show the genome-wide distribution of per-region median pathogenic scores of all non-coding SNVs associated to a given protein-coding gene. Two sets of non-coding SNVs are represented: n=4’960’178 common SNVs without clinical assertions (collectively associated to 18196 protein-coding genes; in grey) and n=737 high-confidence non-coding pathogenic variants (collectively associated to 282 monogenic Mendelian disease genes; in orange). Six types of genomic regions are depicted: intergenic, intronic, 3’UTR, 5’UTR, upstream and downstream regions of associated genes. The downstream region is however not represented in the case of pathogenic non-coding SNVs due to the low set size. Five pathogenicity scores are represented: **A.** CADD non-coding score ^9^; **B.** DeepSEA functional significance score ^13^; **C.** Eigen score ^15^; **D.** FunSeq2 score ^11^; and **E.** ReMM scores ^19^. The horizontal red line is depicted in each panel at the median value of the 5’UTR distribution for common non-coding SNVs. **Table S3** reports the two-sided Wilcoxon test p-values evaluating the null hypothesis that the median pathogenicity score distribution in 5’UTR for common non-coding SNVs is not lower than the corresponding distribution for pathogenic variants in the 5 types of genomic regions evaluated, i.e: intergenic, intronic, 3’UTR, 5’UTR and upstream.

**Figure S3.**
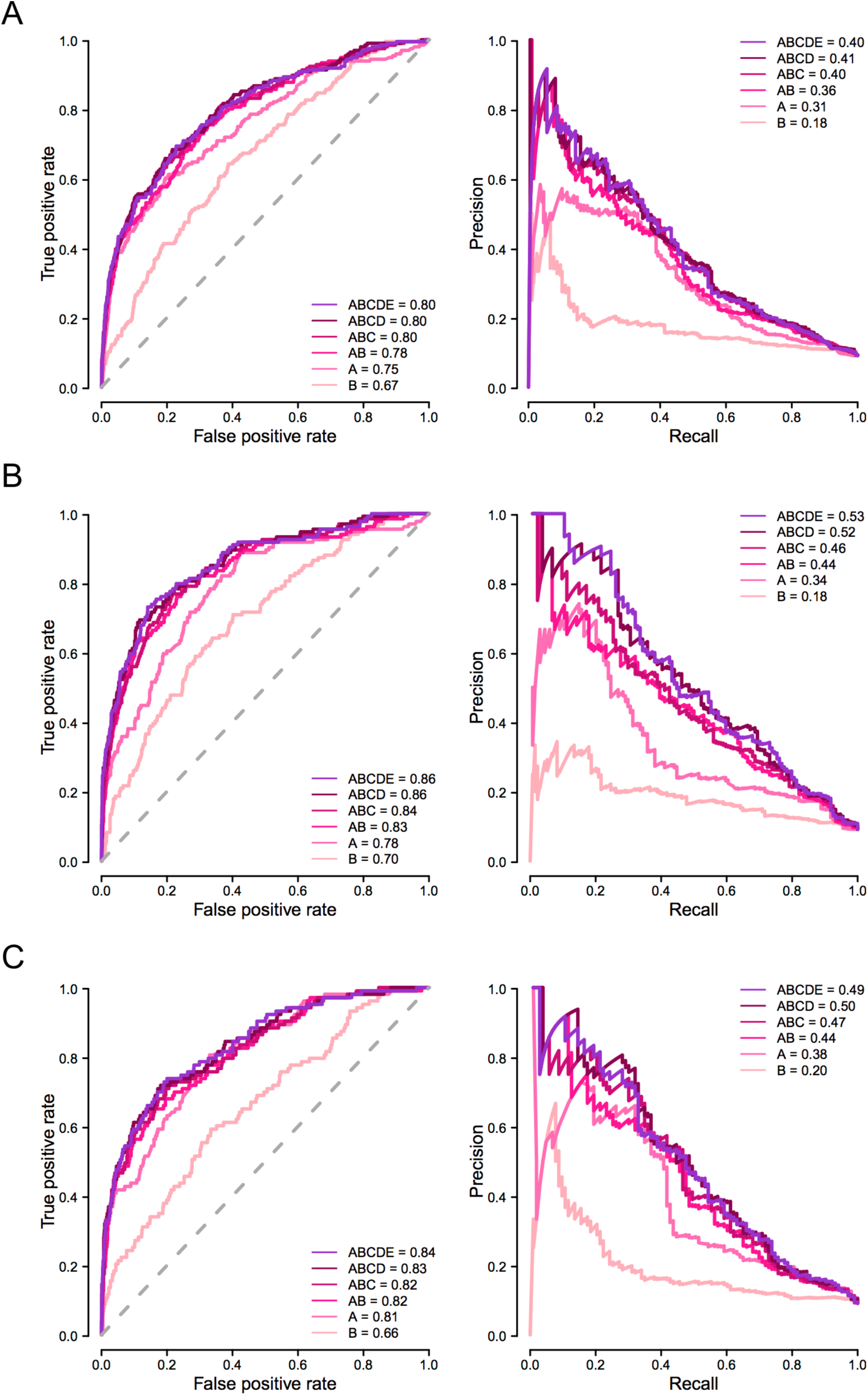
Comparative performance of NCboost models trained upon different sets of features analyzed independently for each source of pathogenic variants. The figure represents the area under the receiver operating characteristic curve (AUROC; **Left panels**) and the area under the Precision-Recall curve (AUPRC; **Right panels**) obtained for each of the six feature configurations evaluated (feature categories A, B, A+B, A+B+C, A+B+C+D and A+B+C+D+E) when trained and tested mimicking a ten-fold cross-validation on high-confidence pathogenic non-coding SNVs from the HGMD-DM set (**Panel A**), Clinvar (**Panel B**) and Smedley’2016 (**Panel C**).

**Figure S4.**
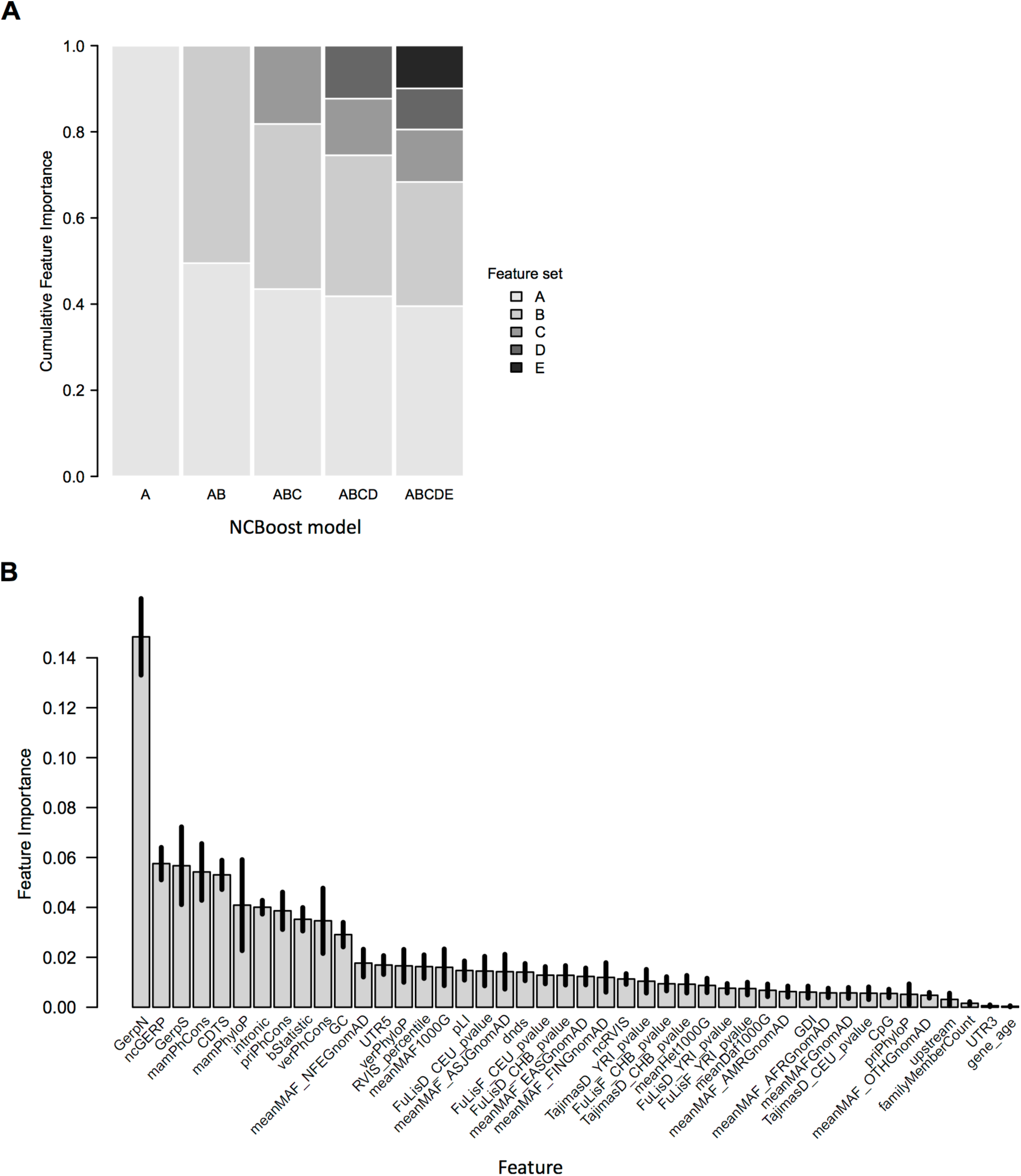
Features importance analysis of NCBoost. **A.** Cumulative feature importance of the feature categories A-E under the six NCBoost feature configurations (A, B, A+B, A+B+C, A+B+C+D and A+B+C+D+E) when trained on n=283 high-confidence pathogenic non-coding SNVs and n=2830 common variants without clinical assertions corresponding to the model performances represented in **Figure 3**. Cumulative importance values were averaged across the 10 independently trained models within the NCBoost bundle, consecutively excluding in each of them 1 of the 10-genome partitions as described in **Methods**. **B**. Mean and standard deviation of individual feature importance values across the 10 independently trained models within the NCBoost bundle (feature configuration ABCD), consecutively excluding in each of them 1 of the 10-genome partitions (**Methods**).

**Figure S5.**
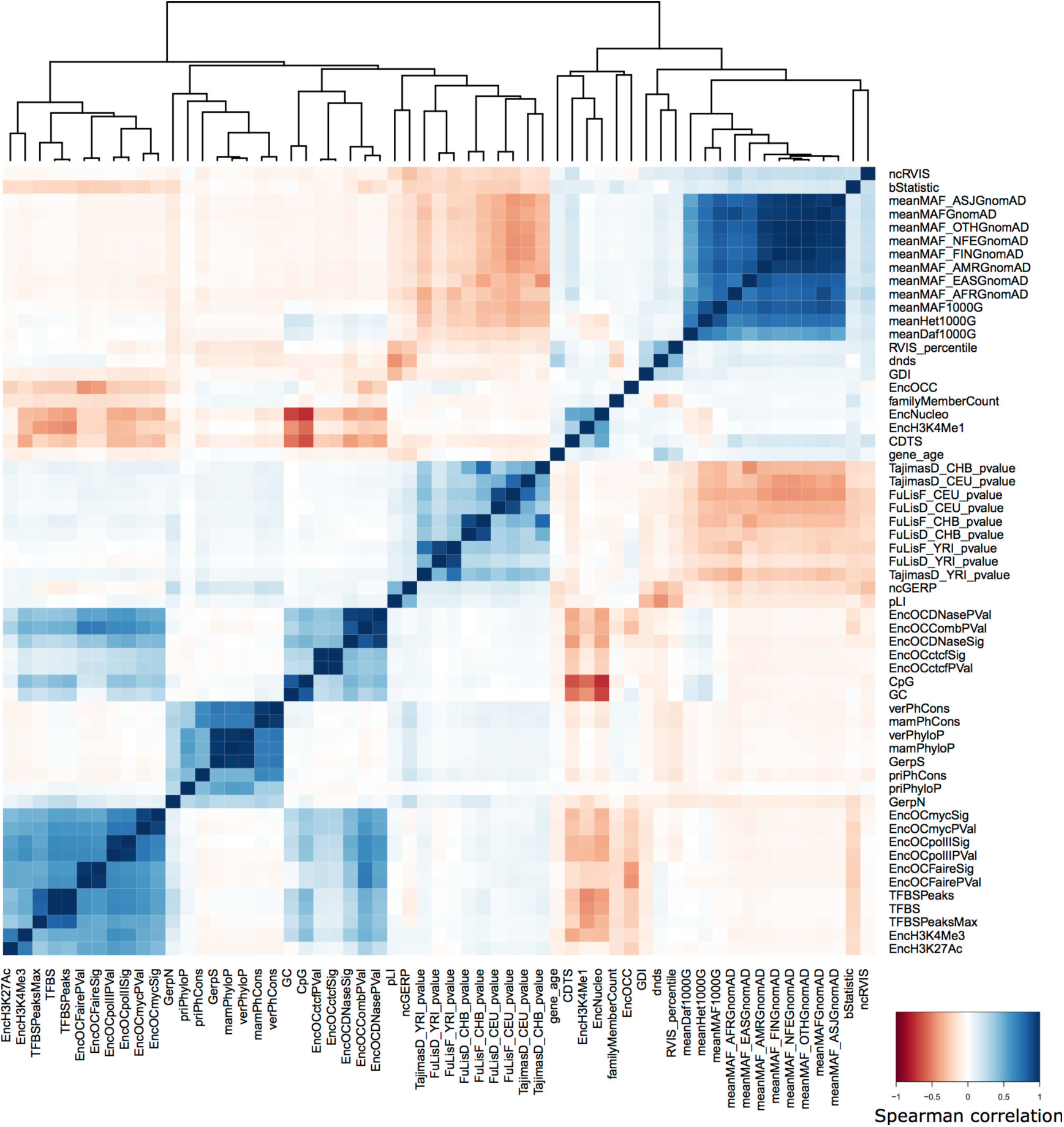
Correlation structure among the features mined in the work. The figure shows the heatmap representation and associated hierarchical clustering of features based on their Spearman correlation values across the a set of SNVs composed of n=737 non-coding pathogenic variants associated to monogenic mendelian disease genes and n=7370 random common variants.

**Figure S6.**
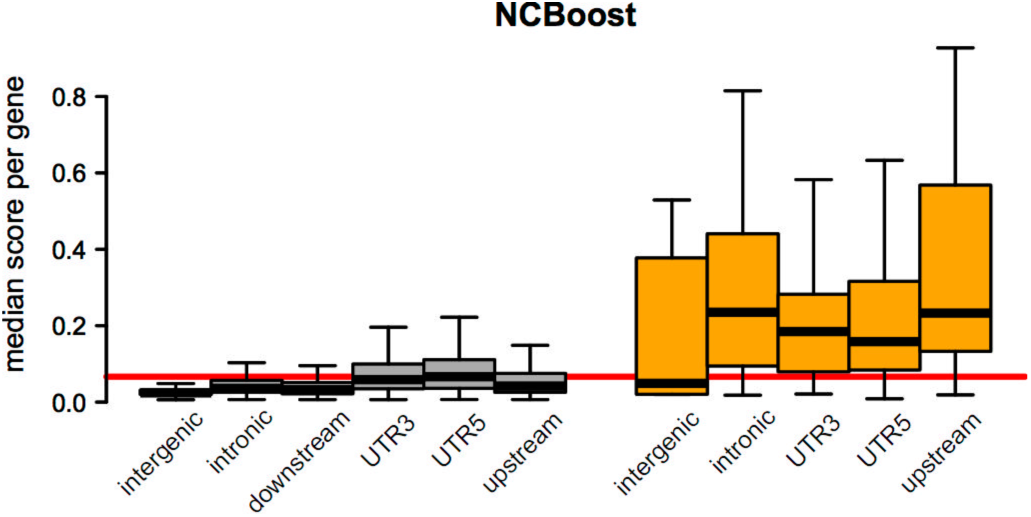
Distribution of NCBoost scores of non-coding SNVs according to the affected type of genomic region. The figure represents the analogous distributions represented in Figure S2, this time for NCBoost scores. Boxplots in the panels show the genome-wide distribution of per-region median pathogenic scores of all non-coding SNVs associated to a given protein-coding gene. Two sets of non-coding SNVs are represented: n=4’960’178 common SNVs without clinical assertions (collectively associated to 18196 protein-coding genes; in grey) and n=737 high-confidence non-coding pathogenic variants (collectively associated to 282 monogenic Mendelian disease genes; in orange). Six types of genomic regions are depicted: intergenic, intronic, 3’UTR, 5’UTR, upstream and downstream regions of associated genes. The downstream region is however not represented in the case of pathogenic non-coding SNVs due to the low set size. **Table S3** reports the two-sided Wilcoxon test p-values evaluating the null hypothesis that the median pathogenicity score distribution in 5’UTR for common non-coding SNVs is not lower than the corresponding distribution for pathogenic variants in the 5 types of genomic regions evaluated, i.e: intergenic, intronic, 3’UTR, 5’UTR and upstream.

**Figure S7:**
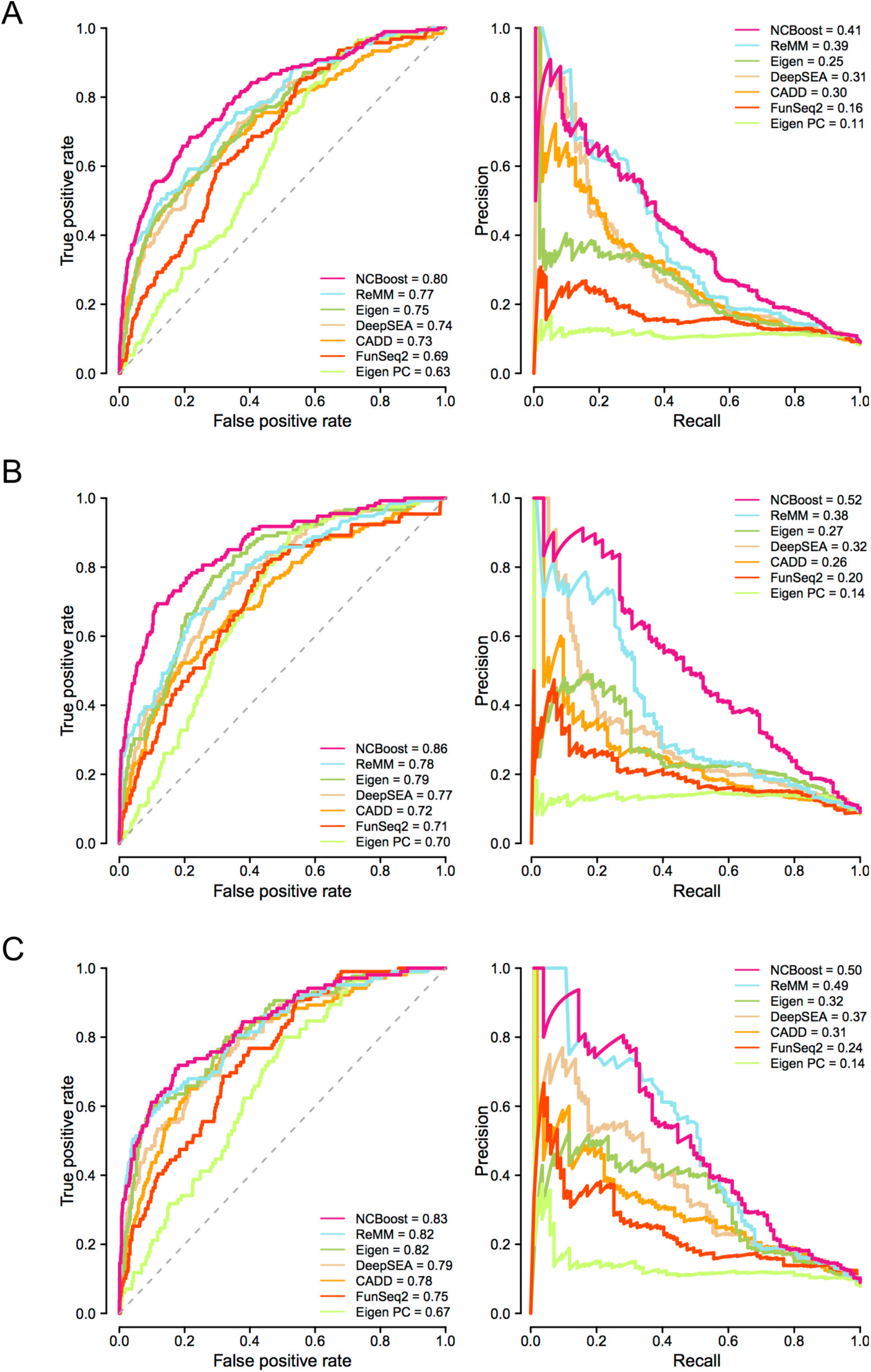
Comparative performance of NCBoost against state-of-the-art methods analyzed independently for each source of pathogenic variants. Figure shows the the area under AUROC (**Left panels**) and the AUPRC (**Right panels**) obtained for NCBoost (feature configuration ABCD) together with 6 state-of-the-art methods (CADD, DeepSEA, Eigen, Eigen-PC, FunSeq2 and ReMM; **Methods**) when tested on high-confidence pathogenic non-coding SNVs from the HGMD-DM set (**Panel A**), Clinvar (**Panel B**) and Smedley’2016 (**Panel C**). The NCBoost model used in each panel as well as the corresponding ‘positive’ and ‘negative’ variants correspond to those described in **Figure 4** for analogous panels.

**Figure S8.**
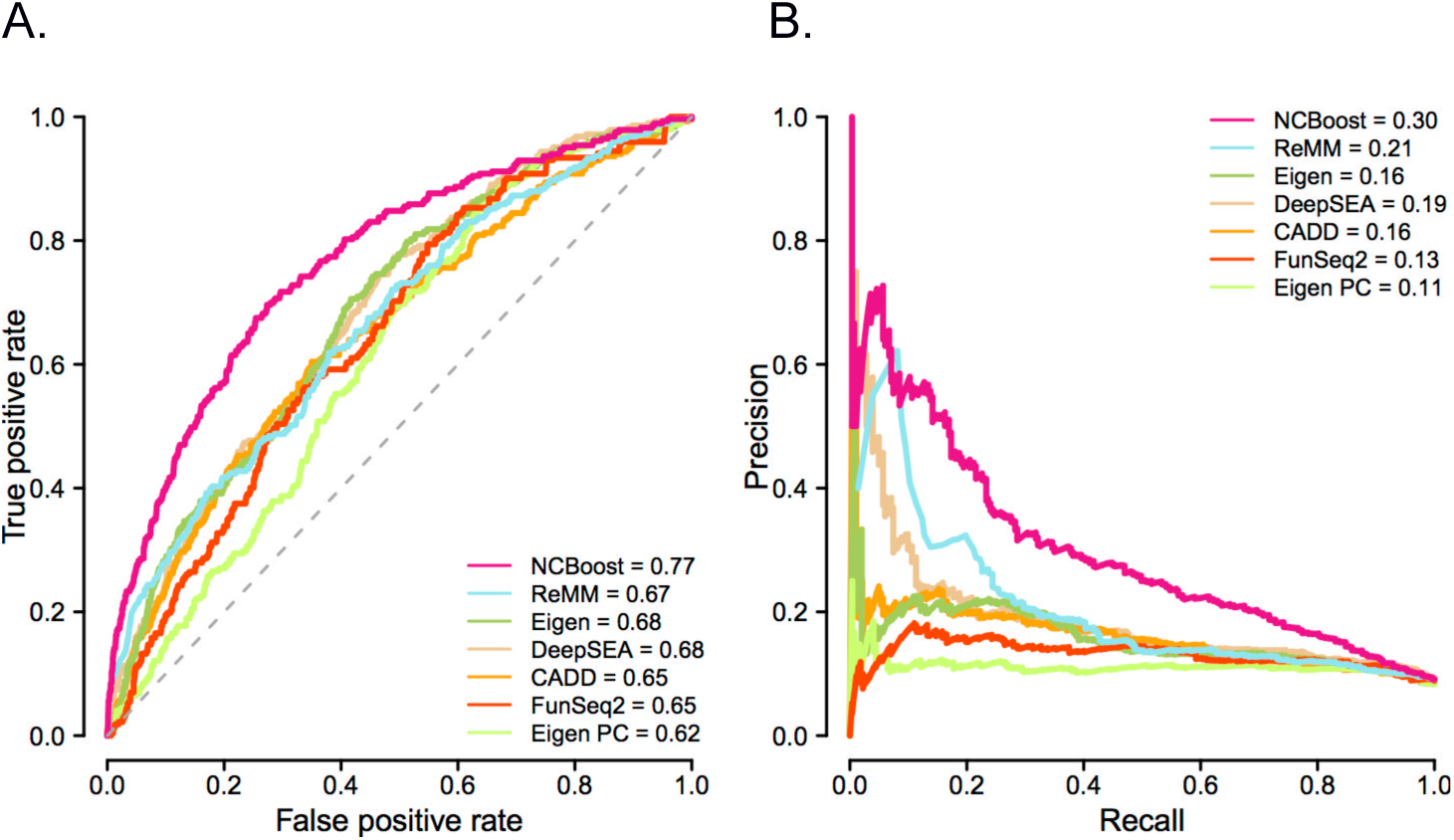
NCBoost capacity to discriminate pathogenic non-coding SNVs from randomly selected rare variants. Figure shows the represents the area under AUROC (**Panel A**) and the AUPRC (**Panel B**) obtained for NCBoost (configuration of features ABCD) together with 6 state-of-the-art methods (CADD, DeepSEA, Eigen, Eigen-PC, FunSeq2 and ReMM; **Methods**) when tested on the same ‘positive set’ of n=283 high-confident set of pathogenic non-coding SNVs as in **Figure 4** and on a negative set that -rather than of common variants-is composed of 2830 randomly selected rare variants (allele frequency < 1%) matched by region. We note that no re-training of NCBoost was done here but used the same NCBoost_ABCD_ model trained as described for Figure 3 and Figure 4.

**Figure S9.**
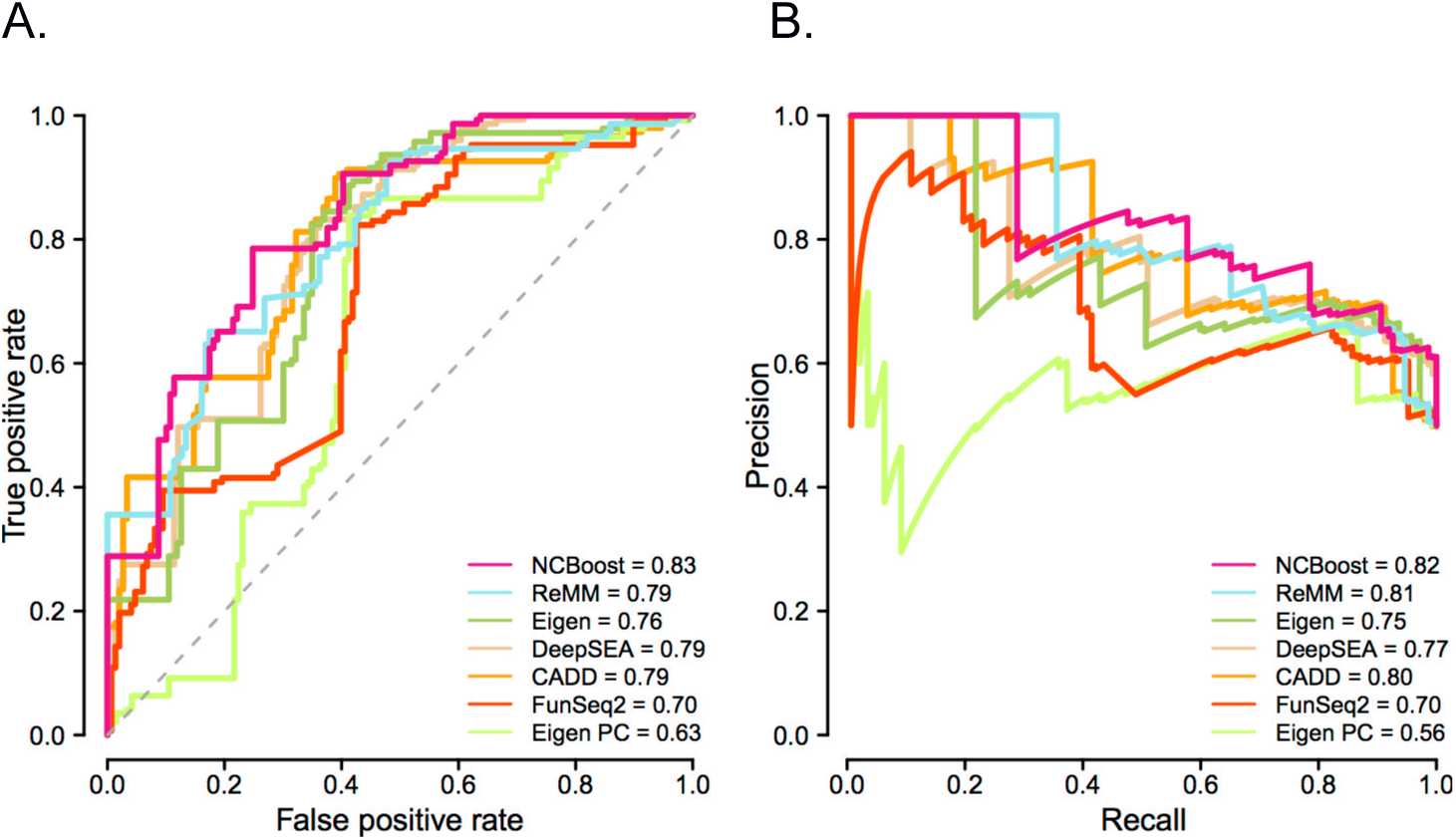
NCBoost capacity to discriminate pathogenic and non-pathogenic variants within the same non-coding region of a given gene. Figure shows the represents the area under AUROC (**Panel A**) and the AUPRC (**Panel B**) obtained for NCBoost (configuration of features ABCD) together with 6 state-of-the-art methods (CADD, DeepSEA, Eigen, Eigen-PC, FunSeq2 and ReMM; **Methods**) when tested on a set of 149 region-matched pairs of pathogenic and random common variants associated to 54 unique genes. We note that no re-training of NCBoost was done here but used the same NCBoost_ABCD_ model trained as described for Figure 3 and Figure 4.

## References

1. McKusick-Nathans Institute of Genetic Medicine, Johns Hopkins University (Baltimore, MD) (2018). Online Mendelian Inheritance in Man, OMIM^®^, https://omim.org/.

2. Institute of Medicine (US) Committee on Accelerating Rare Diseases Research and Orphan Product Development (2010). 2, Profile of Rare Diseases. In Rare Diseases and Orphan Products: Accelerating Research and Development, Field, MJ, and Boat, TF, eds. (Washington (DC): National Academies Press (US)), p. https://www.ncbi.nlm.nih.gov/books/NBK56184/.

3. Chong, J.X., Buckingham, K.J., Jhangiani, S.N., Boehm, C., Sobreira, N., Smith, J.D., Harrell, T.M., McMillin, M.J., Wiszniewski, W., Gambin, T., et al. (2015). The Genetic Basis of Mendelian Phenotypes: Discoveries, Challenges, and Opportunities. Am. J. Hum. Genet. 97, 199–215.

4. Wright, C.F., FitzPatrick, D.R., and Firth, H.V. (2018). Paediatric genomics: diagnosing rare disease in children. Nat. Rev. Genet. 19, 253–268.

5. Zappala, Z., and Montgomery, S.B. (2016). Non-Coding Loss-of-Function Variation in Human Genomes. Hum. Hered. 81, 78–87.

6. Hindorff, L.A., Sethupathy, P., Junkins, H.A., Ramos, E.M., Mehta, J.P., Collins, F.S., and Manolio, T.A. (2009). Potential etiologic and functional implications of genome-wide association loci for human diseases and traits. Proc. Natl. Acad. Sci. 106, 9362–9367.

7. Farh, K.K.-H., Marson, A., Zhu, J., Kleinewietfeld, M., Housley, W.J., Beik, S., Shoresh, N., Whitton, H., Ryan, R.J.H., Shishkin, A.A., et al. (2015). Genetic and epigenetic fine mapping of causal autoimmune disease variants. Nature 518, 337–343.

8. Telenti, A., Pierce, L.C.T., Biggs, W.H., di Iulio, J., Wong, E.H.M., Fabani, M.M., Kirkness, E.F., Moustafa, A., Shah, N., Xie, C., et al. (2016). Deep sequencing of 10,000 human genomes. Proc. Natl. Acad. Sci. 113, 11901–11906.

9. Kircher, M., Witten, D.M., Jain, P., O’Roak, B.J., Cooper, G.M., and Shendure, J. (2014). A general framework for estimating the relative pathogenicity of human genetic variants. Nat. Genet. 46, 310–315.

10. Ritchie, G.R.S., Dunham, I., Zeggini, E., and Flicek, P. (2014). Functional annotation of noncoding sequence variants. Nat. Methods 11, 294–296.

11. Fu, Y., Liu, Z., Lou, S., Bedford, J., Mu, X.J., Yip, K.Y., Khurana, E., and Gerstein, M. (2014). FunSeq2: a framework for prioritizing noncoding regulatory variants in cancer. Genome Biol. 15, 480.

12. Lee, D., Gorkin, D.U., Baker, M., Strober, B.J., Asoni, A.L., McCallion, A.S., and Beer, M.A. (2015). A method to predict the impact of regulatory variants from DNA sequence. Nat. Genet. 47, 955–961.

13. Zhou, J., and Troyanskaya, O.G. (2015). Predicting effects of noncoding variants with deep learning-based sequence model. Nat. Methods 12, 931–934.

14. Shihab, H.A., Rogers, M.F., Gough, J., Mort, M., Cooper, D.N., Day, I.N.M., Gaunt, T.R., and Campbell, C. (2015). An integrative approach to predicting the functional effects of non-coding and coding sequence variation. Bioinformatics 31, 1536–1543.

15. Ionita-Laza, I., McCallum, K., Xu, B., and Buxbaum, J.D. (2016). A spectral approach integrating functional genomic annotations for coding and noncoding variants. Nat. Genet. 48, 214–220.

16. Dunham, I., Kundaje, A., Aldred, S.F., Collins, P.J., Davis, C.A., Doyle, F., Epstein, C.B., Frietze, S., Harrow, J., Kaul, R., et al. (2012). An integrated encyclopedia of DNA elements in the human genome. Nature 489, 57–74.

17. Kundaje, A., Meuleman, W., Ernst, J., Bilenky, M., Yen, A., Heravi-Moussavi, A., Kheradpour, P., Zhang, Z., Wang, J., Ziller, M.J., et al. (2015). Integrative analysis of 111 reference human epigenomes. Nature 518, 317–330.

18. Stunnenberg, H.G., Hirst, M., Abrignani, S., Adams, D., de Almeida, M., Altucci, L., Amin, V., Amit, I., Antonarakis, S.E., Aparicio, S., et al. (2016). The International Human Epigenome Consortium: A Blueprint for Scientific Collaboration and Discovery. Cell 167, 1145–1149.

19. Smedley, D., Schubach, M., Jacobsen, J.O.B., Köhler, S., Zemojtel, T., Spielmann, M., Jäger, M., Hochheiser, H., Washington, N.L., McMurry, J.A., et al. (2016). A Whole-Genome Analysis Framework for Effective Identification of Pathogenic Regulatory Variants in Mendelian Disease. Am. J. Hum. Genet. 99, 595–606.

20. Ponting, C.P., and Hardison, R.C. (2011). What fraction of the human genome is functional? Genome Res. 21, 1769–1776.

21. Rands, C.M., Meader, S., Ponting, C.P., and Lunter, G. (2014). 8.2% of the Human Genome Is Constrained: Variation in Rates of Turnover across Functional Element Classes in the Human Lineage. PLoS Genet. 10, e1004525.

22. Kellis, M., Wold, B., Snyder, M.P., Bernstein, B.E., Kundaje, A., Marinov, G.K., Ward, L.D., Birney, E., Crawford, G.E., Dekker, J., et al. (2014). Defining functional DNA elements in the human genome. Proc. Natl. Acad. Sci. 111, 6131–6138.

23. Fu, W., and Akey, J.M. (2013). Selection and Adaptation in the Human Genome. Annu. Rev. Genomics Hum. Genet. 14, 467–489.

24. Vitti, J.J., Grossman, S.R., and Sabeti, P.C. (2013). Detecting Natural Selection in Genomic Data. Annu. Rev. Genet. 47, 97–120.

25. Nielsen, R., Hellmann, I., Hubisz, M., Bustamante, C., and Clark, A.G. (2007). Recent and ongoing selection in the human genome. Nat. Rev. Genet. 8, 857–868.

26. King, D.C., Taylor, J., Zhang, Y., Cheng, Y., Lawson, H.A., Martin, J., ENCODE groups for Transcriptional Regulation and Multispecies Sequence Analysis, Chiaromonte, F., Miller, W., and Hardison, R.C. (2007). Finding cis-regulatory elements using comparative genomics: Some lessons from ENCODE data. Genome Res. 17, 775–786.

27. Aggarwala, V., and Voight, B.F. (2016). An expanded sequence context model broadly explains variability in polymorphism levels across the human genome. Nat. Genet. 48, 349–355.

28. Tyekucheva, S., Makova, K.D., Karro, J.E., Hardison, R.C., Miller, W., and Chiaromonte, F. (2008). Human-macaque comparisons illuminate variation in neutral substitution rates. Genome Biol. 9, R76.

29. Taylor, M.S., Massingham, T., Hayashizaki, Y., Carninci, P., Goldman, N., and Semple, C.A.M. (2008). Rapidly evolving human promoter regions. Nat. Genet. 40, 1262-1263-1264.

30. Consortium, T. 1000 G.P. (2010). A map of human genome variation from population-scale sequencing. Nature 467, 1061–1073.

31. Lek, M., Karczewski, K.J., Minikel, E.V., Samocha, K.E., Banks, E., Fennell, T., O’Donnell-Luria, A.H., Ware, J.S., Hill, A.J., Cummings, B.B., et al. (2016). Analysis of protein-coding genetic variation in 60,706 humans. Nature 536, 285–291.

32. di Iulio, J., Bartha, I., Wong, E.H.M., Yu, H.-C., Lavrenko, V., Yang, D., Jung, I., Hicks, M.A., Shah, N., Kirkness, E.F., et al. (2018). The human noncoding genome defined by genetic diversity. Nat. Genet. 50, 333–337.

33. Stenson, P.D., Mort, M., Ball, E.V., Shaw, K., Phillips, A.D., and Cooper, D.N. (2014). The Human Gene Mutation Database: building a comprehensive mutation repository for clinical and molecular genetics, diagnostic testing and personalized genomic medicine. Hum. Genet. 133, 1–9.

34. Landrum, M.J., Lee, J.M., Benson, M., Brown, G., Chao, C., Chitipiralla, S., Gu, B., Hart, J., Hoffman, D., Hoover, J., et al. (2016). ClinVar: public archive of interpretations of clinically relevant variants. Nucleic Acids Res. 44, D862–D868.

35. Wang, K., Li, M., and Hakonarson, H. (2010). ANNOVAR: functional annotation of genetic variants from high-throughput sequencing data. Nucleic Acids Res. 38, e164–e164.

36. Zerbino, D.R., Achuthan, P., Akanni, W., Amode, M.R., Barrell, D., Bhai, J., Billis, K., Cummins, C., Gall, A., Girón, C.G., et al. (2018). Ensembl 2018. Nucleic Acids Res. 46, D754–D761.

37. Siepel, A. (2005). Evolutionarily conserved elements in vertebrate, insect, worm, and yeast genomes. Genome Res. 15, 1034–1050.

38. Hubisz, M.J., Pollard, K.S., and Siepel, A. (2011). PHAST and RPHAST: phylogenetic analysis with space/time models. Brief. Bioinform. 12, 41–51.

39. Pollard, K.S., Hubisz, M.J., Rosenbloom, K.R., and Siepel, A. (2010). Detection of nonneutral substitution rates on mammalian phylogenies. Genome Res. 20, 110–121.

40. Davydov, E.V., Goode, D.L., Sirota, M., Cooper, G.M., Sidow, A., and Batzoglou, S. (2010). Identifying a High Fraction of the Human Genome to be under Selective Constraint Using GERP++. PLoS Comput. Biol. 6, e1001025.

41. Pybus, M., Dall’Olio, G.M., Luisi, P., Uzkudun, M., Carreño-Torres, A., Pavlidis, P., Laayouni, H., Bertranpetit, J., and Engelken, J. (2014). 1000 Genomes Selection Browser 1.0: a genome browser dedicated to signatures of natural selection in modern humans. Nucleic Acids Res. 42, D903–D909.

42. Tajima, F. (1989). Statistical method for testing the neutral mutation hypothesis by DNA polymorphism. Genetics 123, 585–595.

43. Fu, Y.X., and Li, W.H. (1993). Statistical tests of neutrality of mutations. Genetics 133, 693–709.

44. McVicker, G., Gordon, D., Davis, C., and Green, P. (2009). Widespread Genomic Signatures of Natural Selection in Hominid Evolution. PLoS Genet. 5, e1000471.

45. Rausell, A., Mohammadi, P., McLaren, P.J., Bartha, I., Xenarios, I., Fellay, J., and Telenti, A. (2014). Analysis of Stop-Gain and Frameshift Variants in Human Innate Immunity Genes. PLoS Comput Biol 10, e1003757.

46. Itan, Y., Shang, L., Boisson, B., Patin, E., Bolze, A., Moncada-Vélez, M., Scott, E., Ciancanelli, M.J., Lafaille, F.G., Markle, J.G., et al. (2015). The human gene damage index as a gene-level approach to prioritizing exome variants. Proc. Natl. Acad. Sci. 112, 13615–13620.

47. Petrovski, S., Wang, Q., Heinzen, E.L., Allen, A.S., and Goldstein, D.B. (2013). Genic Intolerance to Functional Variation and the Interpretation of Personal Genomes. PLoS Genet 9, e1003709.

48. Petrovski, S., Gussow, A.B., Wang, Q., Halvorsen, M., Han, Y., Weir, W.H., Allen, A.S., and Goldstein, D.B. (2015). The Intolerance of Regulatory Sequence to Genetic Variation Predicts Gene Dosage Sensitivity. PLOS Genet. 11, e1005492.

49. Popadin, K.Y., Gutierrez-Arcelus, M., Lappalainen, T., Buil, A., Steinberg, J., Nikolaev, S.I., Lukowski, S.W., Bazykin, G.A., Seplyarskiy, V.B., Ioannidis, P., et al. (2014). Gene Age Predicts the Strength of Purifying Selection Acting on Gene Expression Variation in Humans. Am. J. Hum. Genet. 95, 660–674.

50. Chen, W.-H., Lu, G., Chen, X., Zhao, X.-M., and Bork, P. (2017). OGEE v2: an update of the online gene essentiality database with special focus on differentially essential genes in human cancer cell lines. Nucleic Acids Res. 45, D940–D944.

51. Consortium, T.E.P. (2012). An integrated encyclopedia of DNA elements in the human genome. Nature 489, 57–74.

52. Chen, T., and Guestrin, C. (2016). XGBoost: A Scalable Tree Boosting System. (ACM Press), pp. 785–794.

53. Friedman, J.H. (2001). Greedy Function Approximation: A Gradient Boosting Machine. Ann. Stat. 29, 1189–1232.

54. Dang, V.T., Kassahn, K.S., Marcos, A.E., and Ragan, M.A. (2008). Identification of human haploinsufficient genes and their genomic proximity to segmental duplications. Eur. J. Hum. Genet. 16, 1350–1357.

55. Quinodoz, M., Royer-Bertrand, B., Cisarova, K., Di Gioia, S.A., Superti-Furga, A., and Rivolta, C. (2017). DOMINO: Using Machine Learning to Predict Genes Associated with Dominant Disorders. Am. J. Hum. Genet. 101, 623–629.

56. Mostafavi, H., Berisa, T., Day, F.R., Perry, J.R.B., Przeworski, M., and Pickrell, J.K. (2017). Identifying genetic variants that affect viability in large cohorts. PLOS Biol. 15, e2002458.

57. Short, P.J., McRae, J.F., Gallone, G., Sifrim, A., Won, H., Geschwind, D.H., Wright, C.F., Firth, H.V., FitzPatrick, D.R., Barrett, J.C., et al. (2018). De novo mutations in regulatory elements in neurodevelopmental disorders. Nature 555, 611–616.

58. Javierre, B.M., Burren, O.S., Wilder, S.P., Kreuzhuber, R., Hill, S.M., Sewitz, S., Cairns, J., Wingett, S.W., Várnai, C., Thiecke, M.J., et al. (2016). Lineage-Specific Genome Architecture Links Enhancers and Non-coding Disease Variants to Target Gene Promoters. Cell 167, 1369–1384.e19.

59. Chen, L., Ge, B., Casale, F.P., Vasquez, L., Kwan, T., Garrido-Martín, D., Watt, S., Yan, Y., Kundu, K., Ecker, S., et al. (2016). Genetic Drivers of Epigenetic and Transcriptional Variation in Human Immune Cells. Cell 167, 1398–1414.e24.

60. Pellacani, D., Bilenky, M., Kannan, N., Heravi-Moussavi, A., Knapp, D.J.H.F., Gakkhar, S., Moksa, M., Carles, A., Moore, R., Mungall, A.J., et al. (2016). Analysis of Normal Human Mammary Epigenomes Reveals Cell-Specific Active Enhancer States and Associated Transcription Factor Networks. Cell Rep. 17, 2060–2074.

61. Schmitt, A.D., Hu, M., Jung, I., Xu, Z., Qiu, Y., Tan, C.L., Li, Y., Lin, S., Lin, Y., Barr, C.L., et al. (2016). A Compendium of Chromatin Contact Maps Reveals Spatially Active Regions in the Human Genome. Cell Rep. 17, 2042–2059.

62. Yuan, X., Song, M., Devine, P., Bruneau, B.G., Scott, I.C., and Wilson, M.D. (2018). Heart enhancers with deeply conserved regulatory activity are established early in development.

63. Smedley, D., and Robinson, P.N. (2015). Phenotype-driven strategies for exome prioritization of human Mendelian disease genes. Genome Med. 7,.

64. Lee, S., Abecasis, G.R., Boehnke, M., and Lin, X. (2014). Rare-Variant Association Analysis: Study Designs and Statistical Tests. Am. J. Hum. Genet. 95, 5–23.

65. He, Z., O’Roak, B.J., Smith, J.D., Wang, G., Hooker, S., Santos-Cortez, R.L.P., Li, B., Kan, M., Krumm, N., Nickerson, D.A., et al. (2014). Rare-variant extensions of the transmission disequilibrium test: application to autism exome sequence data. Am. J. Hum. Genet. 94, 33–46.

66. Pybus, M., Dall’Olio, G.M., Luisi, P., Uzkudun, M., Carreño-Torres, A., Pavlidis, P., Laayouni, H., Bertranpetit, J., and Engelken, J. (2014). 1000 Genomes Selection Browser 1.0: a genome browser dedicated to signatures of natural selection in modern humans. Nucleic Acids Res. 42, D903–D909.

